# Insight into Canary Island pine physiology provided by stable isotope patterns of water and plant tissues along an altitudinal gradient

**DOI:** 10.1101/2021.03.30.437734

**Authors:** José Carlos Miranda, Marco M. Lehmann, Matthias Saurer, Jan Altman, Kerstin Treydte

**Author notes:** Author of correspondence: Jose Carlos Miranda.

## Abstract

The Canary Islands, an archipelago east of Morocco’s Atlantic coast, present steep altitudinal gradients covering various climatic zones from hot deserts to subalpine Mediterranean, passing through fog-influenced cloud forests. Unlike the majority of the Canarian flora, *Pinus canariensis* grow along most of these gradients, allowing the study of plant functioning in contrasting ecosystems. Here we assess the water sources (precipitation, fog) of *P. canariensis* and its physiological behavior in its different natural environments. We analyzed carbon and oxygen isotope ratios of water and organics from atmosphere, soil and different plant organs and tissues (including 10-year annual time series of tree-ring cellulose) of six sites from 480 to 1990 m asl on the Canary island La Palma. We found a decreasing δ^18^O trend in source water that was overridden by an increasing δ^18^O trend in needle water, leaf assimilates and tree-ring cellulose with increasing altitude, suggesting site-specific tree physiological responses to relative humidity. Fog-influenced and fog-free sites showed similar δ^13^C values, suggesting photosynthetic activity to be limited by stomatal closure and irradiance at certain periods. Besides, we observed an ^18^O-depletion (fog-free and timberline sites) and ^13^C-depletion (fog-influenced and fog-free sites) in latewood compared to earlywood caused by seasonal differences in: (i) water uptake (i.e. deeper ground water during summer drought, fog water frequency and interception) and (ii) meteorological conditions (stem radial growth and latewood δ^18^O correlated with winter precipitation). In addition, we found evidence for foliar water uptake and strong isotopic gradients along the pine needle axis in water and assimilates. These gradients are likely the reason for an unexpected underestimation of pine needle water δ^18^O when applying standard leaf water δ^18^O models. Our results indicate that soil water availability and air humidity conditions are the main drivers of the physiological behavior of pine along the Canary Island’s altitudinal gradients.

## Introduction

The Canary Archipelago is a natural laboratory for the study of plant physiology and plant responses to climate, due to its strong climate variability (dry, temperate, continental and polar climates; AEMET-IM 2012) across relatively small space (averaging 1000 km^2^ per island). This is caused by the mixed effect of the wide altitudinal range (0 to 3718 m asl) and the influence of the humid trade winds, and triggers an altitudinal-belt structured flora (Aguilar et al. 2010). Northeastern trade winds influence the windward slopes by increasing air humidity, causing orographic precipitation and leading to fog formation (more frequent in summer) at mid altitudes (∼1250 m asl), while leeward slopes face drier conditions (Máyer and Marzol 2013). Fog increases vegetation water availability by means of fog interception that in optimal conditions can reach three times the amount of precipitation water (Aboal et al. 2013; Marzol and Máyer 2012). Interestingly, the main landscape-forming tree species Canary Island pine (*Pinus canariensis* C. Sm. ex DC. in Buch) with an European paleodistribution (Miranda 2017), spreads over several altitudinal climate-driven flora belts, growing from 200 to 2200 m asl on both windward and leeward slopes, with precipitation ranging from 300 to 3000 mm/year and annual average temperatures from 10 to 20 °C (Climent et al. 1996; Pérez-de-Paz et al. 1994). Thus, Canary Island pine is an ideal candidate to study tree physiological responses to climate variability and species-specific strategies to survive under a wide climatic range. Previous studies on this species revealed species-specific strategies to cope with drought (López et al. 2013), frost (Luis et al. 2007), volcanic eruptions (López de Heredia et al. 2014; Miranda et al. 2020; Rodríguez Martín et al. 2013) and fires (Rozas et al. 2013), demonstrating its plasticity. Those studies were, however, restricted to short time periods or single locations, and therefore have not assessed at once the full ecological spectrum of the species along altitudinal gradients so far, leaving open how the physiology of the species adapts to such a wide range of climatic conditions.

Carbon (δ^13^C) and oxygen (δ^18^O) stable isotope ratios in plant tissue, like leaves and tree rings, can be used as a proxy to assess both plant physiology and plant water relations (Farquhar et al. 2007; Gessler et al. 2014; McCarroll and Loader 2004). δ^18^O of plant tissue depends on meteorological, physical, physiological and morphological processes and variables at leaf level (Brinkmann et al. 2018; Cernusak et al. 2016; Farquhar et al. 2007; Tang and Feng 2001). In leaves, water inputs from diverse sources, which can have different isotope composition, occur under dew, fog and rainfall conditions (Burgess and Dawson 2004; Kim and Lee 2011). The isotopic signature of the leaf water is transferred on leaf assimilates (Lehmann et al. 2017). In the phloem, newly produced sugars mix with older carbohydrate reserves dampening the isotopic signature of the sugar pool (McCarroll and Loader 2004; Sternberg and Deniro 1983). Finally, during tree-ring cellulose biosynthesis oxygen atom exchange occurs between phloem sugars and ^18^O-depleted xylem water (Sternberg et al. 2006).

Conversely, δ^13^C in tree samples is determined by the intracellular CO_2_ concentration during leaf carbohydrate synthesis, which is under strong control of stomatal conductance and photosynthetic rate and therefore reflecting intrinsic water use efficiency (Gessler et al. 2014; McCarroll and Loader 2004), with both processes being regulated by environmental and species-specific conditions. After photosynthesis, further isotope fractionations can occur at the leaf and stem levels together with mixing of sugar pools with different assimilation times or storage (Gessler and Treydte 2016), causing generally a basipetal ^13^C enrichment in phloem sugars (Boegelein et al. 2019; Brueggemann et al. 2011; Offermann et al. 2011). Although the mentioned isotope fractionations and mixing processes occur during the vertical transport of sugars, tree-ring cellulose δ^13^C and δ^18^O values can be strongly correlated with the isotopic signature of leaf sugars and reflect physiological processes at the leaf level, specifically in evergreen species (Gessler et al. 2014).

Thus, δ^13^C and δ^18^O values of plant compounds can be a useful tool to assess plant responses to altitude-driven environmental variability. The carbon isotopic composition in plants is known to become generally ^13^C-enriched with increasing altitude due to decreasing temperatures, and CO_2_ and O_2_ atmospheric partial pressures (Körner et al. 1991), but this could be modified by concurrent changes in humidity. Variations of δ^18^O values of tree tissues in relation to altitude are not so well studied. Plant compounds are thought to be ^18^O-depleted with increasing altitude following the isotopic patterns of meteoric water (Barbour 2007; Burk and Stuiver 1981; Naoe et al. 2016; Terwilliger et al. 2002). However, under strong influence on leaf water ^18^O enrichment, opposite trends have also been reported (Kahmen et al. 2011). Both tree-ring δ^13^C and δ^18^O also respond to seasonal environmental or source water variations (Gessler et al. 2014; Treydte et al. 2014) and hence, to seasonal physiological variations.

Besides, stands influenced by recurrent seasonal fog events can present inter and intra-annual variation in δ^13^C and δ^18^O due to isotopically differentiated source water inputs (Prada et al. 2015, Johnstone et al. 2013) and due to changes in temperature, relative humidity and irradiation affecting isotope fractionation processes (Ritter et al. 2019). Trees can intercept fog water, this is, small fog droplets impact on their leaves, branches and stems, and merge into bigger drops (Ritter et al. 2008). Needle-like leaves are better fog droplet interceptors than other leaf morphologies or bigger vegetative surfaces (Shuttleworth 1977), and the capacity of Canary Island pine to intercept large amounts of fog water (2 times the precipitation water input) has been already assessed (Aboal et al. 2000; Aboal et al. 2013). The intercepted fog drops increase tree water availability by either flowing down and dripping to the ground where roots can take it up, or by directly transferring into the leaves as observed for several cloud forest species (Goldsmith et al. 2013).

Here we aim at understanding Canary Island pine physiology and its relation to hydroclimatic conditions as a function of altitude. For this goal, we investigate the influence of altitude-driven climate and its seasonal variations on δ^13^C and δ^18^O of twigs, needles, and early- and latewood cellulose of pine trees and on the isotopic signature of the source water on the island of La Palma. Sampling was performed at five sites along an altitudinal gradient on the windward slope (fog influence at mid-altitudes) and at one site on the leeward slope (fog-free). We hypothesize (i) a site-specific strategy to avoid water losses (higher water use efficiency at drier sites triggered by lower transpiration rates), allowing the species to grow in a wide range of relative humidity conditions. This would imply higher δ^13^C and δ^18^O values in plant tissues with increasing altitude according to Scheidegger’s et al (2000) conceptual model. We also hypothesize (ii) seasonal differences in average stomatal conductance at locations where summer drought is not alleviated by fog influence; and in photosynthetic rates at locations where frequent fog events occur seasonally. This would mean differences in both δ^13^C and δ^18^O in earlywood (formed in spring and early summer) compared to latewood (formed in summer and autumn). Finally, we hypothesize (iii) an influence of fog intercepted water/foliar water uptake on tree rings from the mid-altitude site and, thus, the capability of this species’ needles to take up water. This would mean an ^18^O enrichment, as fog water was reported to be enriched compared to precipitation water (Prada et al. 2015).

## Materials and Methods

### Site description

The study was performed on La Palma (Canary Islands) on Canary Island pine (*Pinus canariensis* C. Sm. ex DC. in Buch) along an altitudinal gradient ranging from 480 to 1990 m asl. Climate of the Canary Islands is strongly modulated by altitude and exposure (windward and leeward slopes). Under stable atmospheric conditions, northeastern humid trade winds affect mostly mid-altitude locations typically between 1200-1300 m asl, exceptionally extending to 800-1800 m asl (Dorta 1985; Marzol 2008; Torres et al. 2001; Valladares 1995). Above the trade wind influence, a subsidence temperature inversion avoids water vapor to rise up, causing its condensation and resulting in fog conditions - enabling fog water interception by vegetation and even foliar water uptake - and orographic precipitation at mid altitudes (Herrera et al. 2001). Thus, precipitation (mainly occurring during the wet season, from October to March) increases from coastal to mid altitudes and decreases above the trade wind zone (Fig. 1), where it is completely absent from June to August (Máyer and Marzol 2013). Temperatures usually decrease with increasing elevation (Fig. 1), except during subsidence temperature inversion periods (Bechtel 2016).

**Figure 1:**
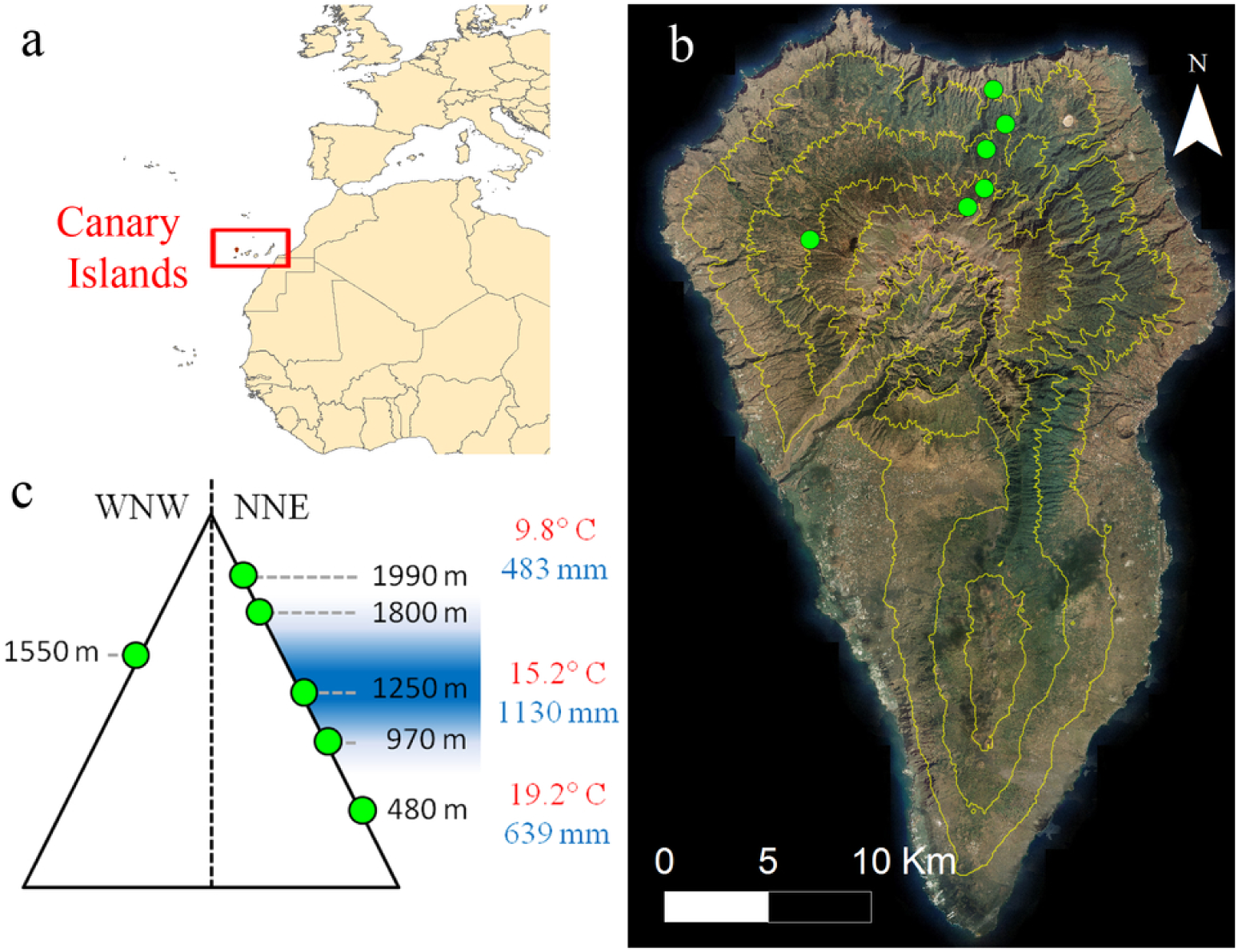
Canary Islands and study sites on La Palma. a) Location of the Canary Islands, b) La Palma island, contour lines (500 m) and selected study sites (green dots), c) schematic diagram of site locations along the altitudinal gradient, aspect, fog influence (blue color) and annual average temperatures and precipitation at low, mid and high altitudes.

Five sites were selected along an altitudinal transect on an NNE slope at the following altitudes (Fig. 1): 480 m (28° 49’ 25.16” N, 17° 50’ 37.12” W), 970 m (28° 48’ 31.16” N, 17° 50’ 13.86” W), 1250 m (28° 47’ 50.80” N, 17° 50’ 47.42”W), 1800 m (28° 46’ 48.71” N, 17° 50’ 48.98”W) and 1990 m (28° 46’ 19.39”N, 17° 51’ 18.07”W). Trade wind and hence, fog influence, is strongest at the mid-altitude site (fog influenced, 1250 m), rare at the 1800 and 970 m sites, and absent at the timberline (1990 m) and the lowest, 480 m site. A sixth site was assessed on the WNW slope (leeward) at 1550 m (28° 45’ 21.94” N, 17° 55’ 57.26”) that is not influenced by the trade winds, holding as a fog-free control in the altitudinal range where fog commonly occurs on the windward slope.

### Sampling of plant tissues, soil and water

Samples from different pine organs and tissues, as well as soil and atmospheric water vapor samples, were taken to perform δ^18^O, δ^13^C and dendrochronological analyses at two sampling days between 10 am and 3 pm (Eidg. Forschungsanstalt WSL 2018). During the first sampling day on the 7^th^ of March 2018, soil, stem, twig and needle material were sampled from five trees at each of the five NNE slope sites. Soil samples (S) were collected at 10 cm depth. In each tree, a stem xylem piece (SX) of 5-8 cm length was collected with a 0.5 cm increment borer (Haglof, Sweden). A 1×8 cm piece of stem phloem (SP) was cut using a chisel. Two twigs of five to eight cm length from the lower part of the canopy were collected with a pruner, and their bark (TB) and xylem (TX) were separated and collected together with previous growing season needles from 2017 (Ne). Samples were placed in gas-tight 12 ml glass vials (“Exetainer”, Labco, Lampeter, UK), stored in cooling bags with ice packs and then frozen after the field work.

During the second sampling day on the 9^th^ of March 2018, additional needle samples from previous growing season (2017) were taken at the 1250 and 1990 m sites, divided in three equal parts from the brachyblast to the tip (i.e. “bottom”, “mid”, and “tip”) and collected separately in Exetainers (full length of the needles varied from 15 to 30 cm). During the second sampling day, also water vapor (WV) was sampled in cryogenic traps at the 480 m, 1250 m and 1990 m sites at two consecutive times (10:30 am −12:30 pm and 12:30-2:30 pm). U-glass traps were connected to a 3 mm tube on one side (inlet) and a small air pump (1 L/min) on the other side (outlet) and submerged into a dry ice/ethanol slurry in dewar flasks in order to condensate all water vapor flowing through the glass trap for 2 h.

Later during the fieldtrip (second week of March 2018), four cores per tree were sampled from 30 trees at the fog-influenced 1250 m and timberline 1990 m sites (NNE slope) and at the fog-free 1550 m site (WNW slope) with 0.5 cm diameter increment borers for tree-ring analyses (tree-ring width, earlywood and latewood δ^13^C and δ^18^O).

### Sample processing and stable isotope analyses

Water from soil, stem xylem and phloem, twig xylem and bark, and needles was cryogenically extracted according to West et al. (2006): Sample tubes were placed into a 80°C water bath, connected to a vacuum system including water traps cooled with liquid nitrogen. Captured water was then pipetted to 2 ml vials. δ^18^O values of all these samples and water vapor samples were determined with a high-temperature conversion/elemental analyzer coupled via a ConFlo III referencing interface to a Delta^Plus^XP isotope ratio mass spectrometer (TC/EA-IRMS, all supplied by Finnigan MAT, Bremen, Germany), using a glassy carbon reduction method as previously described (Gehre et al. 2004). Oxygen isotope ratios were measured on the resulting gas CO and referenced to VSMOW (Vienna Standard Mean Ocean Water, IAEA, Vienna, Austria). Measurement precision of quality control standards were <0.2 ‰ (standard deviation).

All dry needle material from the sites at 1250 and 1990 m was ground to a fine powder using a steel ball-mill. Aliquots (1 mg) of needle organic matter (Ne-OM) were weighted in silver capsules. Needle water-soluble compounds (Ne-WSC), which mostly but not exclusively reflect the photosynthetic assimilates, were extracted from 100 mg needle organic matter in a 2 ml vial in a water bath at 85 °C for 30 min using 1.5 ml deionized water (Lehmann et al. 2015). Subsequently, samples were centrifuged (2 min, 10,000 *g*) and the supernatant contacting the Ne-WSC was transferred to a new vial. Aliquots (c. 1 mg) were then pipetted to silver capsules, frozen at −20°C, and freeze-dried. Both δ^13^C and δ^18^O of Ne-OM and Ne-WSC were measured by TC/EA-IRMS (vario PYRO cube, Elementar, Hanau, Germany; coupled to the same Conflo and IRMS as for water analysis) and referenced to VPDB and VSMOW, respectively. Laboratory control standard was typically measured with a precision (standard deviation) of <0.3 ‰ for δ^13^C and 0.2 ‰ for δ^18^O.

For dendrochronological analyses, the surface of all sampled cores was cut with a microtome (Gartner and Nievergelt 2010), and polished afterwards with a 600 grained sandpaper to maximize visibility of the tree rings. Tree-ring widths were measured with 0.01 mm resolution with a LINTAB measuring device (Rinntech, Heidelberg, Germany) and cross-dated (Stokes and Smiley 1996) visually and statistically with the software COFECHA (Holmes 1983). Standard dendrochronological procedures were followed to accurately date the tree rings. In order to analyze intra-seasonal tree-ring δ^13^C and δ^18^O variations, three different trees per site and one core per tree were selected (fog-influenced at 1250 m, fog-free at 1550 m and timberline at 1990 m). We avoided cores with distinct compression wood, though it has been reported recently for other conifer species that compression wood only has a minor if any effect on the stable isotope signature of tree ring cellulose (Janecka et al. 2020). Earlywood (EW) and latewood (LW) of each ring for the period 2008-2017 was split off with a scalpel under a stereomicroscope. EW and LW samples were transferred to Teflon filter bags (Ankom Technology, Macedon, NY, USA) and holocellulose was extracted using a two-step digestion method (NaOH and NaClO_2_) following Boettger et al. (2007). Extracted cellulose was homogenized following Laumer et al. (2009) and weighted (1 mg) into silver capsules for δ^18^O and δ^13^C analyses as described above.

### Stomatal conductance measurements

During the first sampling day (7^th^ of March 2018), stomatal conductance (g_s_), and leaf temperature measurements were performed 4 times in 2017 needles on each of the studied trees at the 1250 m, 1800 m, and 1990 m sites on the same twigs collected for isotope analyses using an SC-1 leaf porometer (Decagon Devices, Washington, USA).

### Foliar water uptake (FWU)

In order to test the foliar water uptake capacity of Canary Island pine, 0.8-1 m long branches were taken in the field on the 7^th^ of March 2018 at 1250 m and 1990 m asl from the same five study trees used for the previously described tree tissues’ sampling. Branches were placed in black plastic bags to avoid transpiration and water loss. After the fieldwork, branches were re-cut underwater to 0.4 m length, kept overnight in a bucket and again placed in black plastic bags. On the following day, leaf water potential was measured on two needle bundles of every branch with a Scholander pressure chamber (PMS Instrument Company, Albany, OR, USA) to ensure that branches and needles were fully rehydrated (i.e. water potential was 0 MPa). Subsequently, ten needle bundles from 2017 (3 needles per bundle) per tree were sampled all over the branch plus another ten of the needle bundles from 2016 of the trees at the 1250 m site (needles of 2016 were not always present on the trees at the 1990 m site). We applied the gravimetrical method described in Limm et al. (2009) using a scale (Mettler Toledo Switzerland accuracy 0.01g), submerging the needles in labeled water mixed with tap water (δ^18^O = 55.2 ‰). The bottom of the needle bundle was enclosed by parafilm to avoid water loss and kept over the water surface to avoid water entering the needle through the bundle. Uptake and increase of leaf water content were calculated and corrected for leaf surface water capacity after Limm et al. (2009). Finally, needles were transferred to Exetainers for water extraction.

### Mechanistic modeling

Modeling of evaporative needle water ^18^O enrichment above source water (twig xylem water) was performed using the Craig Gordon (1965) model as modified for application to leaves and given by Farquhar et al. (2007):

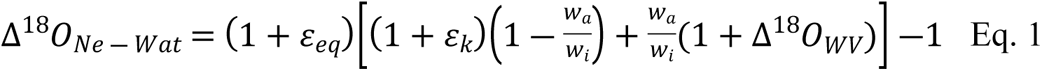

with ε_eq_ and ε_k_ being equilibrium and kinetic fractionation respectively, w_a_/w_i_ the ratio of water vapor partial pressures outside and inside the leaf, and Δ^18^O_WV_ the ^18^O enrichment of water vapor (δ^18^O_WV_)above twig xylem representing the source water (δ^18^O_TX_^)^ (Cernusak et al. 2016):

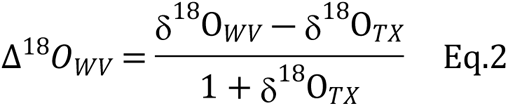

Equilibrium and kinetic fractionation factors as well as water vapor partial pressures were calculated after Cernusak et al. (2016). Air temperature, relative humidity, leaf temperature and g_s_ were measured in situ at 1250, 1800 and 1990 m sites. At 480 and 970 m sites, atmospheric variables were obtained from a station as described in the following section, leaf temperature was approximated to air temperature and g_s_ was approximated to the average g_s_ at the 1250 m site (Table S1). Modeled δ^18^O_Ne-Wat_ values were derived using Eq 2 from Δ^18^O_Ne-Wat_ and δ^18^O_TX_.

To assess the robustness of modeled values, different scenarios were simulated, i.e. varying leaf temperature (measured leaf temperature in each individual ±0.5, ±1 and ±2 standard deviations of the values measured at each site) and relative humidity (average site value ±2.5, ±5 and ±10 %).

In addition, we calculated the apparent biosynthetic fractionation factor (εbio_app_) as the observed δ^18^O differences between the water-soluble compounds and needle water:

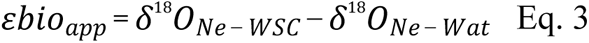

### Meteorological data

To study the influence of climatic variables on tree radial growth and δ^13^C and δ^18^O of tree-ring cellulose, we used temperature and precipitation data from stations at locations with climate conditions comparable to our study sites: One station at 1380 m on a NE slope on La Palma (4.3 km away from our 1250 m site), two stations at 2137 and 2340 on a NE slope on Tenerife (140 km away from 1990 m site), and one station at 1032 m on a WSW slope on Tenerife (130 km away from 1550 m site) (Table 1); the first two stations belong to the Spanish Meteorological Agency (AEMET), and the third one to the local Tenerife Island Government (Cabildo de Tenerife). Relative air humidity at the same stations and further calculation of vapor pressure deficit (VPD) as saturated minus actual vapor pressure, were used to characterize wet and dry seasons at the fog-influenced, fog-free and timberline sites (Table1). Monthly sea-level pressure (SLP) data were taken from the website of the Royal Netherlands Meteorological Institute (Harris et al. 2014).

**Table 1:** Meteorological data of dry and wet seasons for the fog-influenced, fog-free and timberline sites. Mean temperature (Temp), precipitation (Prec), relative humidity (rH) and vapor pressure deficit (VPD) at the tree-ring sites during the dry (April-September) and wet (October-March) seasons over the study period (2008-2017) are given.

| Site | Temp (°C) |  | Prec (mm) |  | rH (%) |  | VPD (kPa) |  |
| --- | --- | --- | --- | --- | --- | --- | --- | --- |
|  | Dry | Wet | Dry | Wet | Dry | Wet | Dry | Wet |
| 1990-timberline | 14.3 | 6.9 | 36.6 | 361.3 | 30.6 | 47.4 | 1.17 | 0.52 |
| 1550 - fog-free | 17.8 | 12.2 | 41.9 | 291.6 | 61.6 | 67 | 0.85 | 0.46 |
| 1250- fog-influenced | 17.32 | 13.78 | 188.1* | 1091.2* | 75.53 | 81.5 | 0.47 | 0.24 |

\* precipitation data does not include water inputs by means of fog water interception that could double or triple annual precipitation

Additionally, sampling day temperatures and relative humidity were measured/estimated. In situ measurements were conducted during the two sampling days at the 1250, 1800 and 1990 m sites with a SC-1 leaf porometer. Both were calculated as the mean of measurements taken immediately before and after the sampling at each site (and after the initial calibration at each site). Meteorological values for the 480 m site at sampling days were gathered from a meteorological station at La Palma (528 m - NE slope, 0.5 km away; AEMET). Values for 970 m site were interpolated from the 480 and the 1250 m sites. (Table 2).

**Table 2:** Meteorological data of the sampling dates for the study sites. Temperature (Temp), relative humidity (rH), vapor pressure deficit (VPD) and stomatal conductance on 2017 needles (*g*_s_) of the two days at the time of water and tissue sampling (water vapor, soil, stem xylem and phloem, twig xylem and bark, and needles) are given, as well as the previous month precipitation decile of the last 30-year February precipitation (Prec_Feb_).

| Date | Sampled tissue | Site | Temp (°C) | rH (%) | VPD (kPa) | $g_s$ (mmol·m<sup>-2</sup>·s<sup>-1</sup>) | Prec<sub>Feb</sub> (decile) |
| --- | --- | --- | --- | --- | --- | --- | --- |
| 07.03.2018 | Water and tissue samples | 1990 m | 11.2<sup>1</sup> | 41<sup>1</sup> | 0.78 | 210.0<sup>+</sup> | 7 |
|  |  | 1800 m | 16.4<sup>1</sup> | 56<sup>1</sup> | 0.82 | 322.7<sup>+</sup> | - |
|  |  | 1250 m | 17.4<sup>1</sup> | 66<sup>1</sup> | 0.68 | 405.9<sup>+</sup> | 9 |
|  |  | 970 m | 18.2<sup>2</sup> | 69<sup>2</sup> | 0.65 | - | - |
|  |  | 480 m | 19.5<sup>3</sup> | 74<sup>3</sup> | 0.59 | - | 8 |
| 09.03.2018 | Needles and water vapor | 1990 m | 9.8 | 74 | 0.31 | - | 7 |
|  |  | 1250 m | 18.3 | 48 | 1.09 | - | 9 |
|  | Water vapor | 480 m | 24.5 | 52 | 1.47 | - | 8 |

<sup>1</sup> Values measured *in situ* with SC-1 leaf porometer
<sup>2</sup> Values interpolated from upper and lower sites.
<sup>3</sup> Values gathered from a meteorological station 500 m away
Other meteorological values collected from stations or calculated as indicated in Materials and methods.
<sup>+</sup> $g_s$ differences in altitude were significant according to the fitted linear mixed model accounting for tree identity as a random factor (P - value < 0.001).

### Statistical analyses

Linear regression models were applied to test for δ^18^O differences among plant tissues, soil and water samples at each site, and among altitudes for each sample type; as well as along needle gradient (bottom, mid and tip parts). The effect of relative humidity at sampling time on needle water δ^18^O was also tested by linear models, as well as the effects of altitude and year on the increase of leaf water content and δ^18^O after submerging the needles. A linear mixed model accounting for tree identity as a random factor was performed to check differences in *g_s_* among altitudes; and accounting for tree identity and year of ring formation as random factors to test for δ^18^O differences in EW or LW, sites, and their interactions. In the case of tree-ring cellulose δ^13^C values, differences between seasonal wood formation were tested independently at each site with a linear mixed model accounting for tree identity and year of ring formation as random factors. Finally, linear models testing the effects of mean annual relative humidity and VPD on δ^18^O of tree-ring cellulose were performed. The non-parametric Kruskal-Wallis test was applied when data did not meet assumptions of normality and homoscedasticity of residuals. This was the case for δ^18^O of SX and SP among altitudes, ring widths among sites and tree-ring δ^13^C among sites for each season of wood formation. To test for differences between groups, Tukey and Dunn post-hoc tests were performed in the case of parametric (linear regression and linear mixed models) and non-parametric (Kruskal-Wallis) analyses, respectively. To determine whether the increase in leaf water content was significantly above 0 after submerging the needles, a one-sample t-test was applied. The same was also applied to test for significant differences between observed and modeled/simulated δ^18^O values of needle water.

Pearson’s correlation coefficients were calculated between climate data (temperature, precipitation, sea-level pressure and North Atlantic Oscillation) and tree-ring width and EW and LW isotope data, respectively. Correlations were performed with monthly mean climatic values, and with trimester means coinciding with the 3-month drought period for the previous and current growing seasons (September_-1_- October_-1_-November_-1_ - SON_-1_, December_-1_-January-February - DJF_-1_, March-April-May - MAM, June-July-August - JJA, September-October-November - SON).

Statistical analyses were carried out in R version 3.6.0 (R Core Team 2019) with the package nlme (Pinheiro et al. 2020) to perform linear mixed models. Figures were produced using the package ggplot2 (Wickham 2009).

## Results

### δ^18^O of soil, water and plant tissues along the altitudinal gradient

δ^18^O values in water decreased significantly (P-value < 0.001) and linearly with increasing altitude for soil (−1.7 ‰ km^-1^), twig xylem (−1.6 ‰ km^-1^) and twig bark (−1.4 ‰ km^-1^) and not in a linear way for stem xylem (from −3.1 to −5.2 ‰) and stem phloem (from −3.0 to −5.5 ‰,), with no statistically significant differences within each site (Fig. 2, Table 3). In contrast, δ^18^O values in needle water increased with altitude from 5.4 to 21.4 ‰ (P-value < 0.001), in an exponential way, with relatively little change at the lower sites. δ^18^O values in needle organic matter and water-soluble compounds increased with increasing altitude and were higher compared to needle water: 17.0 and 20.2 ‰ higher at the 1250 site and 12.6 ‰ and 14.5 ‰ higher at the 1990 m site, respectively. Besides, Ne-WSC were ^18^O-enriched compared to the 10-year average of tree-ring cellulose, by 1.9 ‰ at the 1250 site and 1.7 ‰ at the 1990 site. δ^18^O values of water vapor averaged −16.8 ‰, but did not change significantly along altitude (P-value > 0.1). Relative humidity at sampling time significantly influenced δ^18^O values in needle water (decreasing δ^18^O with increasing humidity) explaining 87.3 % of its variability (P-value < 0.001).

**Table 3:**
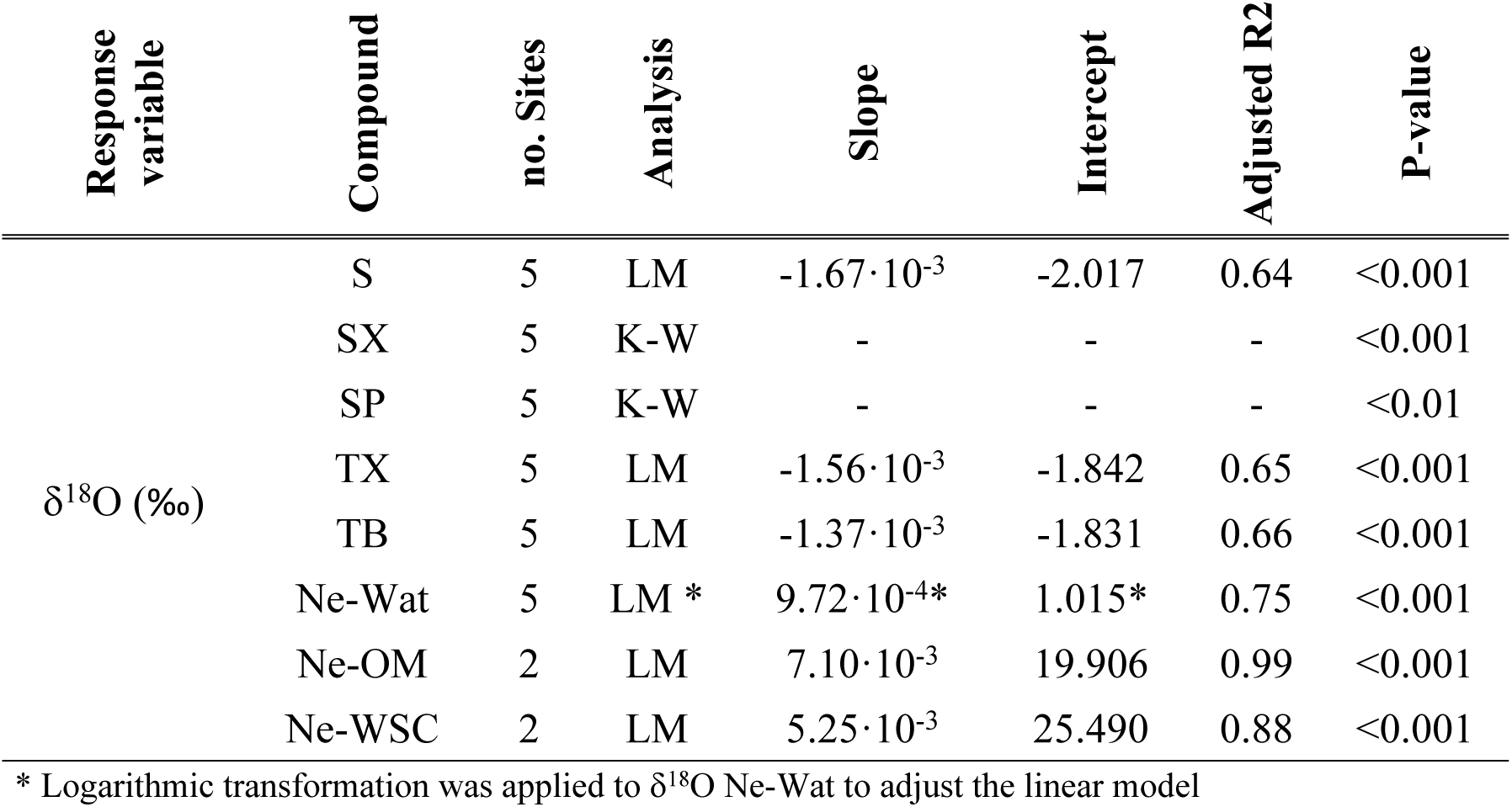
Statistics for the analysis of altitudinal effects on the oxygen isotopic composition of water and assimilates of the different tree tissues and organs. Slope and intercept of the applied linear model are given. LM: linear model, K-W: Kruskal-Wallis. Compounds: soil water (S), stem xylem water (SX), stem phloem water (SP), twig xylem water (TX), twig bark water (TB), needle water (Ne-Wat), needle organic matter (Ne-OM) and water-soluble compounds from needles (Ne-WSC).

Stomatal conductance (*g*_s_) significantly decreased along the altitudinal gradient (P-value < 0.001) averaging 405.9, 322.7 and 210.0 mmol·m^-2^·s^-1^ at 1250 m, 1800 m and 1990 m sites respectively (Table 2). Meteorological conditions at the first sampling day represented a typical wet season day, preceded by heavy rains during the previous weeks. On the second sampling day, however, atmospheric instability caused an inversion in humidity (Table 1 and 2).

**Figure 2:**
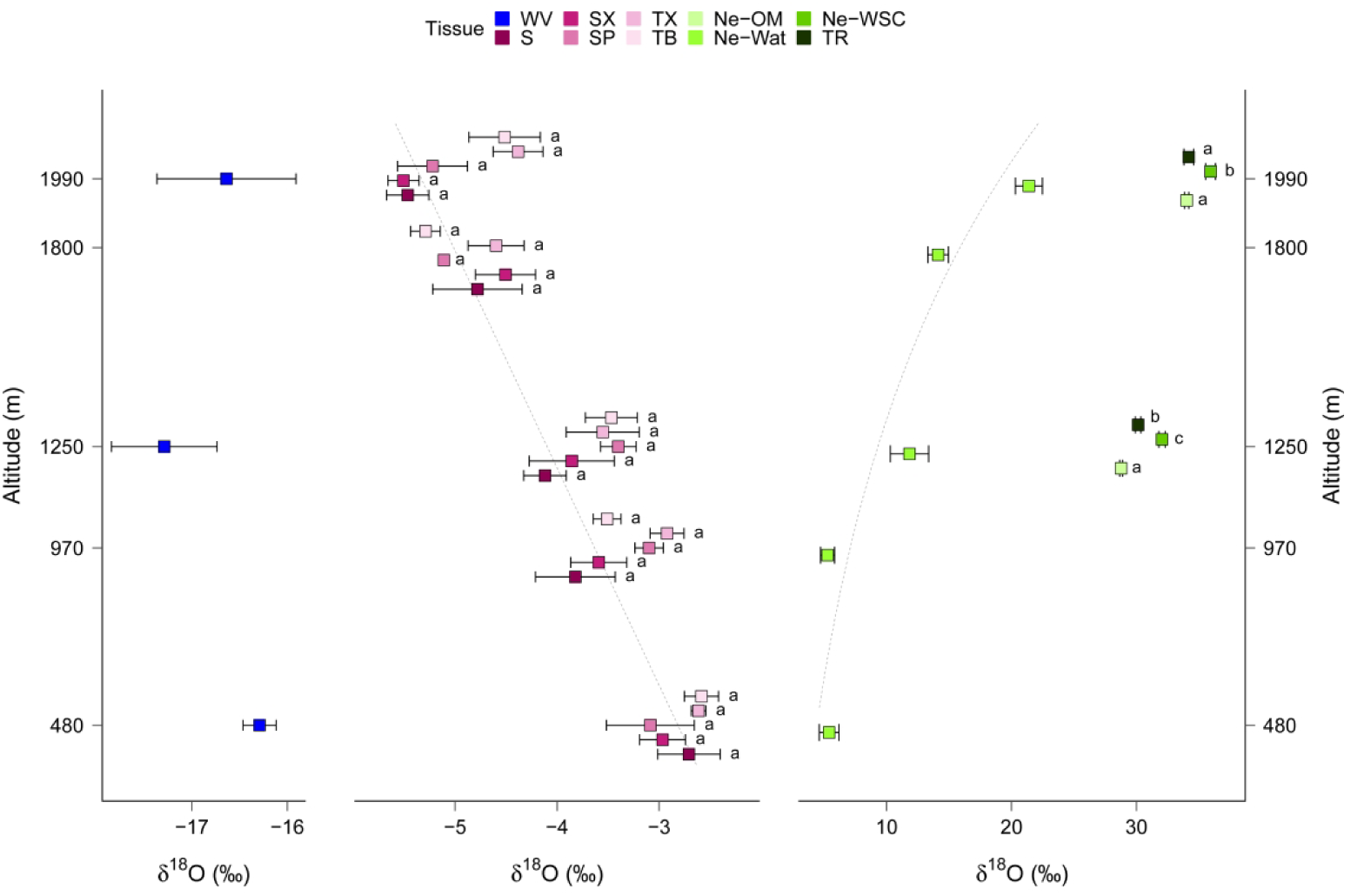
Oxygen isotope composition (δ^18^O) of atmospheric water and extracted water of soil and Canary Island pine samples along an altitudinal gradient on La Palma. Abbreviation: water vapor (WV), soil water (S), stem xylem water (SX), stem phloem water (SP), twig xylem water (TX), twig bark water (TB), needle water (Ne-Wat), needle organic matter (Ne-OM), needle water-soluble compounds (Ne-WSC), and 10-year average of tree-ring cellulose (TR). Dotted lines show linear model calculated for δ^18^O values (left) of soil water and log(δ^18^O) values of Ne-Wat (right). Letters indicate statistically significant differences between tissues at each altitude (Tukey test P < 0.05). Mean values ± standard error (WV n = 2; TR n = 3, other compounds n = 5).

### Needle water δ^18^O: Gradients, mechanistic modeling and FWU

Needle water and Ne-WSC showed a significant (P-value < 0.001 for both compounds) ^18^O enrichment along the needle length (∼15 to 30 cm) from the bottom to the tip at both study sites. δ^18^O values of needle water were 17.0 ‰ higher at the tip than at the bottom of needles at 1250 m, but only 5.3 ‰ higher at 1990 m. In contrast, the δ^18^O range of Ne-WSC along the needles was lower (7.2 ‰) at 1250 m than at 1990 m (11.5 ‰; see differences between Ne-WSC and needle water in Fig. 3). Accordingly, the apparent biosynthetic fractionation factor (ebio_app_) showed opposed trends at the two altitudes. Values of ebio_app_ decreased from the bottom to the tip (25.2 to 15.4 ‰) at 1250 m, but increased from 30.0 to 36.1 ‰ at 1990 m.

**Figure 3:**
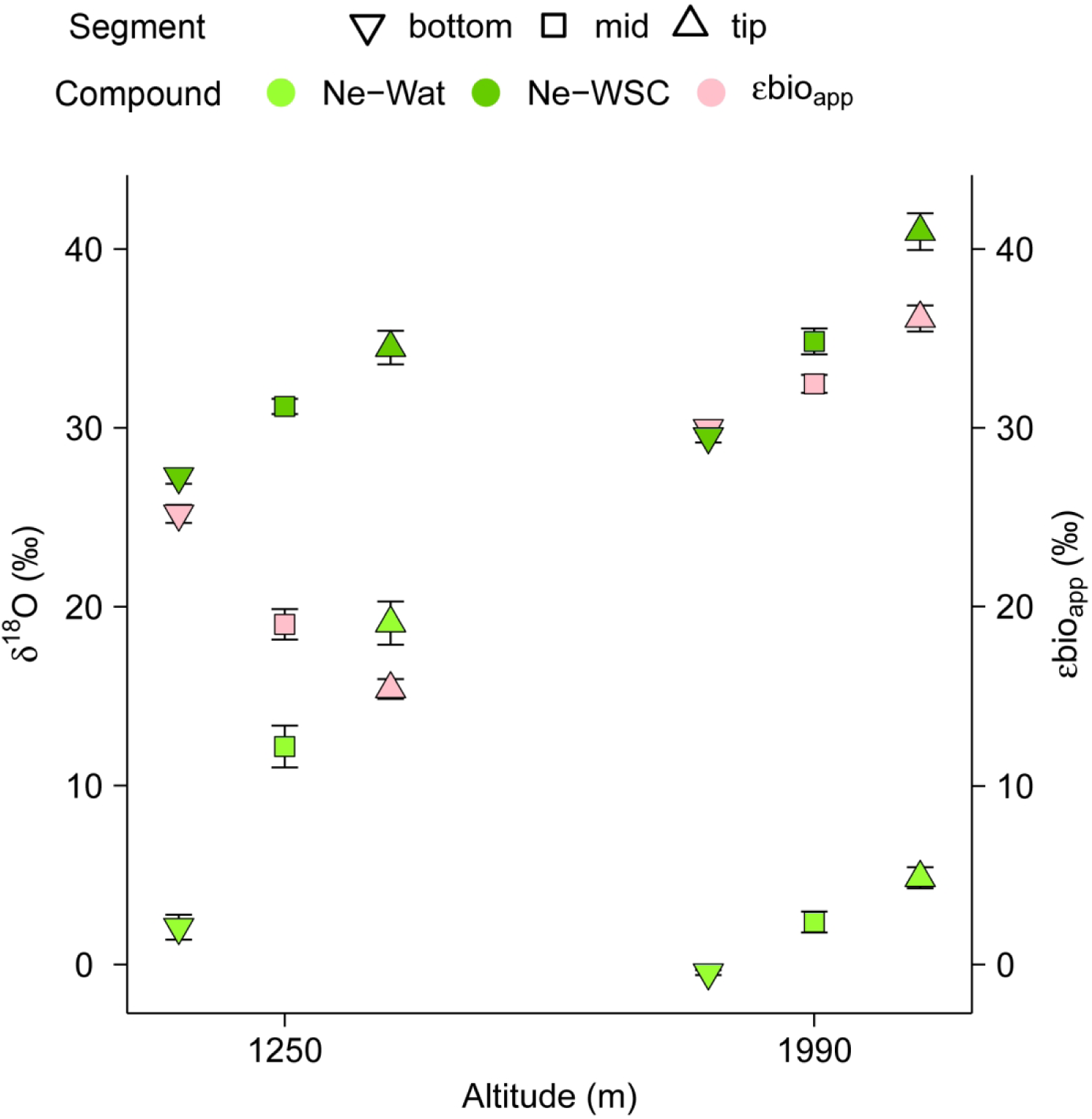
Oxygen isotope composition (δ^18^O) of Canary Island pine needle water and assimilates from the two sites at 1250 and 1990 m asl. δ^18^O values of needle water (Ne-Wat), needle water-soluble compounds (Ne-WSC), and the biosynthetic fractionation factor (εbio_app_) of three needle segments (bottom, mid, tip) are given. Needles ranged from 15 to 30 cm in length. Mean values ± standard error (n = 5).

Modeled δ^18^O values of needle water at the study sites based on Eq. 1 fitted with the observed values, except for those at 1800 m (Fig. 4). Besides, there was a higher variability in observed values compared to modeled ones (average standard deviation of 2.19 in observed values vs. 1.26 in modeled ones). Nevertheless, modeled and observed values were significantly correlated (r = 0.88, P-value < 0.01). The simulations varying leaf temperature input, showed higher dispersions at 480, 970 and 1250 m sites compared to 1800 and 1990 m sites (Fig. S1). Compared to these simulations, dispersions due to relative humidity sensitivity analysis were similar or lower. At 1800 m site, observed values where significantly lower than simulated values (P-value < 0.05), except for the higher leaf temperature scenario (+ 2 standard deviations) and the - 5 and −10% relative humidity scenarios.

**Figure 4:**
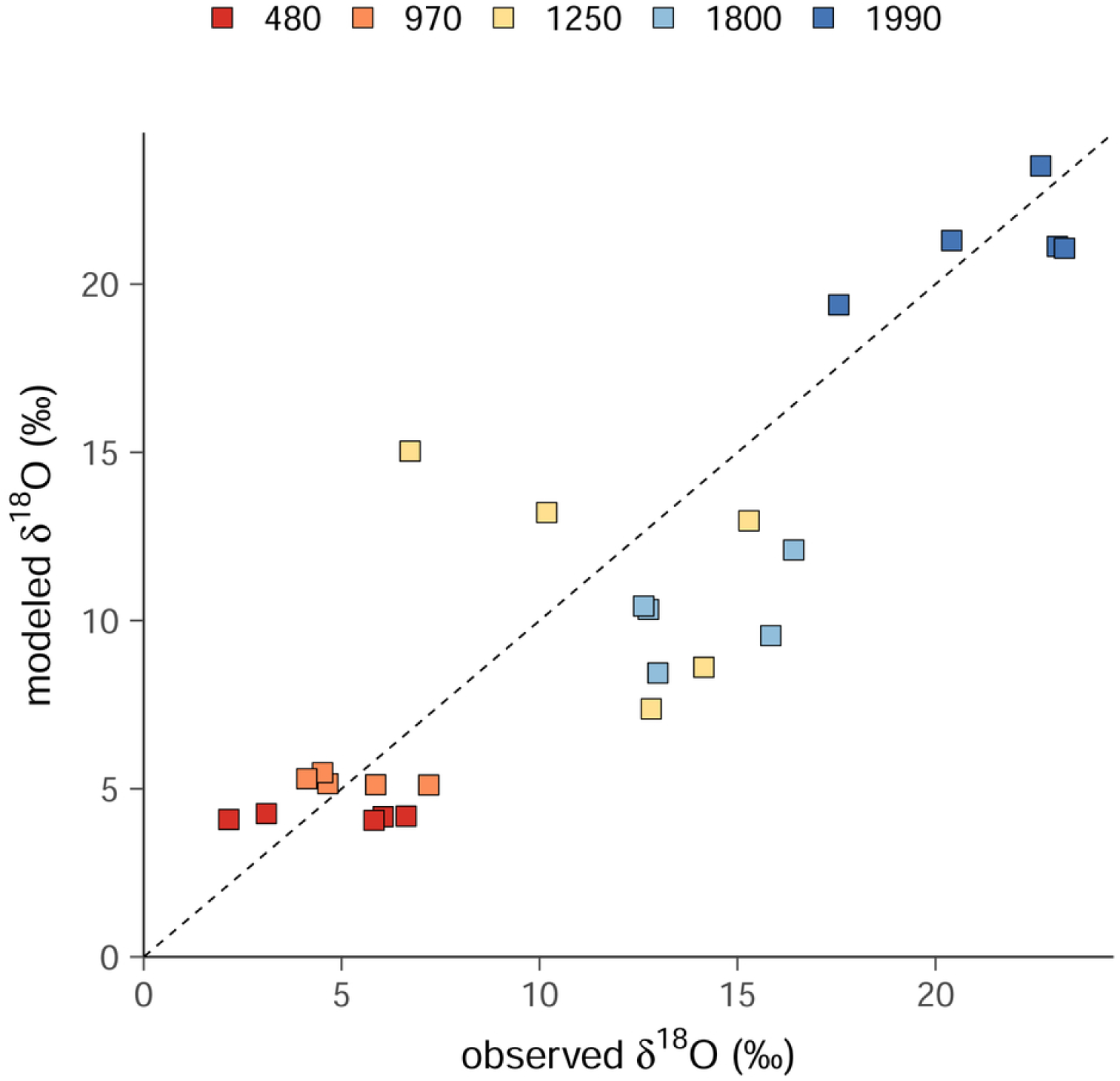
Observed vs. modeled δ^18^O values in needle water of Canary Island pine along an altitudinal gradient on La Palma. Black dotted line represents the 1:1 line.

The capacity of Canary Island pine needles for foliar water uptake (FWU) was demonstrated with needles from two years (2016 and 2017) and two sites (1250 and 1990 m), as the increase in leaf water content (LWC) was significantly above 0 after performing the gravimetric method (P-value < 0.05, Table 4). An increase in LWC > 3 % was observed for the 2017 needles at both sites. 2016 needles (only present at 1990 m) tended to have a 6.4% LWC increase, which was, however, not significant (P-value > 0.05). In addition, δ^18^O values of needle water indicated ^18^O-label uptake after FWU, but showed no significant difference between needle years and altitude (P-value > 0.05).

**Table 4:** Foliar water uptake (FWU) of the studied Canary Island pine needles. Needles from two sites and of two years have been used. ΔLWC: increase in needle water content, δ^18^O_FWU_: δ^18^O values of needle water after foliar water uptake. Mean values ± standard error (n = 5).

| Site | Needle cohort | FWU (gram) | $\Delta\text{LWC}$ (%) | $\delta^{18}\text{O}_{\text{FWU}}$ (‰) |
| --- | --- | --- | --- | --- |
| 1990 m | 2017 | $0.071 \pm 0.014$ | $3.8 \pm 1.9$ | $35.0 \pm 4.5$ |
| 1250 m | 2017 | $0.081 \pm 0.018$ | $3.5 \pm 1.7$ | $33.3 \pm 3.5$ |
| 1250 m | 2016 | $0.124 \pm 0.028$ | $6.4 \pm 1.4$ | $36.3 \pm 5.0$ |

### Tree-ring cellulose δ^18^O and δ^13^C records

Mean δ^18^O values of tree-ring cellulose followed the δ^18^O trend in needle water and assimilates, showing a significant increase along the altitudinal gradient for both EW and LW (Fig. 5a, d). EW increased from 30.3 to 35 ‰ and LW from 29.9 to 33.3 ‰ from the 1250 to the 1990 site. In addition, δ^18^O differences between EW and LW indicated seasonal variations. The δ^18^O values of EW cellulose were significantly higher than LW at the fog-free 1550 m site (0.85 ‰ P-value < 0.001) and timberline 1990 m (1.6 ‰, P-value < 0.001; Fig. 5a, d), while no significant differences were observed at the fog-influenced 1250 m site. Altitudinal-driven environmental changes and seasonal variations in EW vs. LW together explained 82 % of the variability of tree-ring δ^18^O (marginal R^2^ = 0.82).

**Figure 5:**
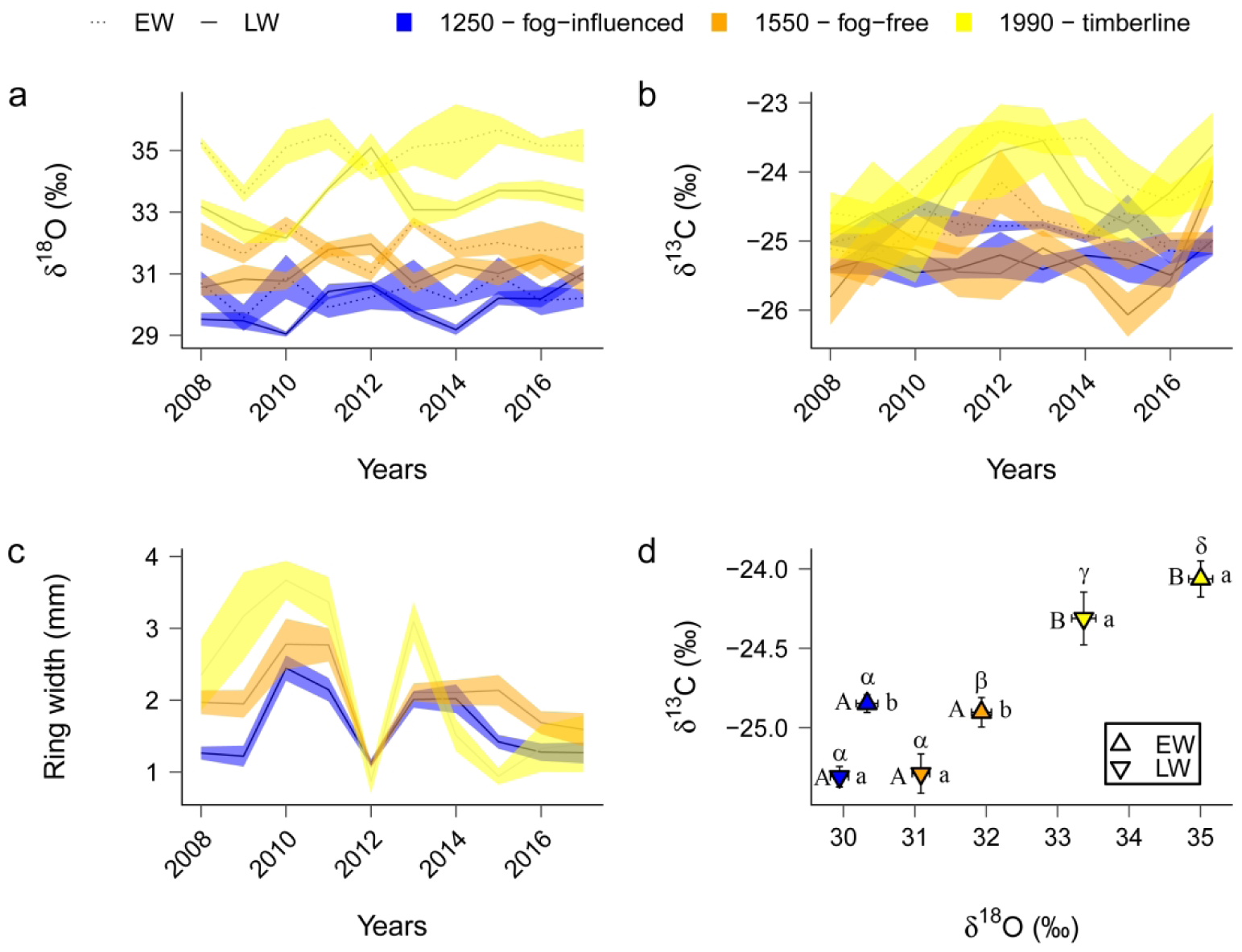
Carbon and oxygen isotope variations of tree-ring cellulose of the Canary Island pines studied on La Palma. Mean a) δ^18^O and b) δ^13^C values of tree-ring cellulose for early (EW) and latewood (LW) of 10-year chronologies (2008-2017) from three sites (fog-influenced at 1250 m, fog-free at 1550 m and timberline at 1990 m). c) Mean ring-width series per year at each site. d) δ^18^O - δ^13^C representation of early- (EW) and latewood (LW) at each site. Greek letters indicate significant differences in δ^18^O values between sites and seasons. Latin letters indicate significant differences in δ^13^C values between seasons at each site (lower case), and between sites for each season (upper case). Both ribbons and error bars show ± standard errors (n = 3).

δ^13^C values of tree-ring cellulose also increased with altitude by 0.81 ‰ for EW and 0.99 ‰ for LW, with higher δ^13^C values at 1990 m compared to both 1250 and 1550 m sites (P-value <0.001 for both EW and LW) (Fig. 5b, d). In contrast to δ^18^O, δ^13^C values of EW were consistently higher compared to LW at 1250 m and 1550 m (P-value < 0.05), but not at 1990 m.

No altitudinal gradient in growth was detected, as tree-ring widths of the 10 years used for isotope analysis were not significantly different among sites. However, tree-ring cellulose δ^18^O increased significantly (P-value < 0.001) with decreasing mean annual relative humidity and with increasing mean annual VPD (Fig. 6). VPD explained 68.5 and 66.4 % of δ^18^O variability in EW and LW cellulose respectively, whereas relative humidity explained 82.2 and 78.5 %.

**Figure 6:**
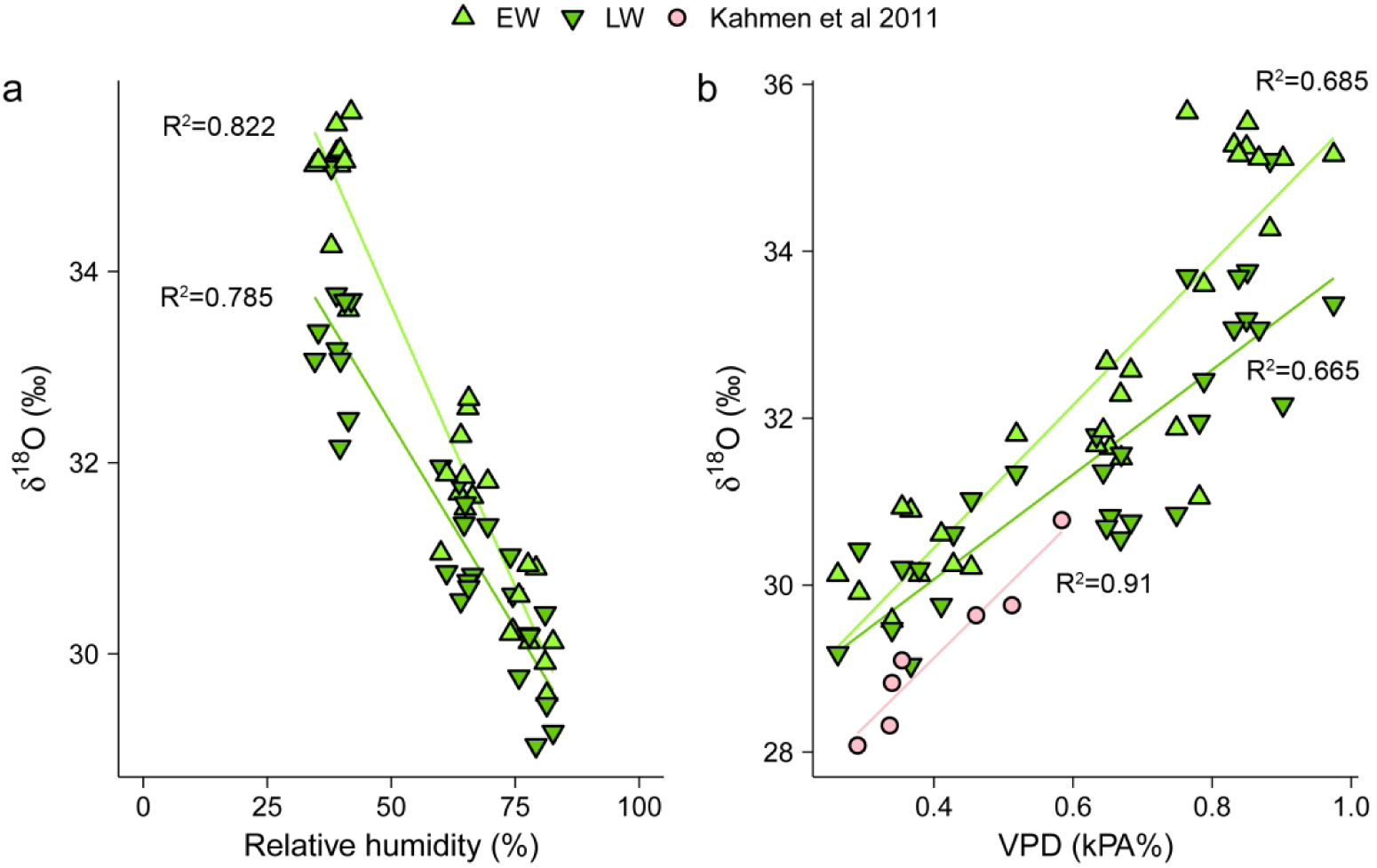
Linear regressions between oxygen isotope values (δ^18^O) of Canary Island pine tree-ring cellulose from earlywood (EW) and latewood (LW) and annual mean relative humidity (left) and vapor pressure deficit (VPD; right), and in comparison to values from Kahmen et al (2011) (right). Points represent mean annual δ^18^O per season (n=3).

We further investigated the climate sensitivity of the tree-ring parameters over the 10-year study period (Fig. 7, S2). Tree-ring width at all sites was positively correlated to previous winter precipitation (DJF_-1_, r between 0.41 and 0.60, P-value < 0.05 for this and following correlations). At the 1550 m fog-free site it was also correlated to spring precipitation (MAM, r = 0.58), and at the 1250 m fog site negatively correlated to autumn precipitation (SON, r = −0.50). LW δ^18^O at 1250 and 1990 m were negatively correlated to previous winter precipitation (DJF_-1,_ r = −0.48 and −0.71, respectively), and also LW δ^13^C at the 1990 m timberline site was negatively correlated to previous winter precipitation (DJF_-1,_ r = −0.38). Regarding sea-level pressure (SLP) negative correlations were found for tree-ring width and previous winter SLP at all sites (DJF_-1_, r between −0.45 and −0.55) and at the 1250 m site also to spring SLP (MAM, r = −0.53). LW δ^18^O was positively correlated to winter SLP at 1250 and 1990 m (r = 0.50 and 0.66 respectively) and at 1250 m also to spring (MAM, r = 0.50) and summer SLP (JJA, r = 0.44). Positive correlation was also found between summer SLP and LW δ^13^C at the fog-free site at 1550 m.

**Figure 7:**
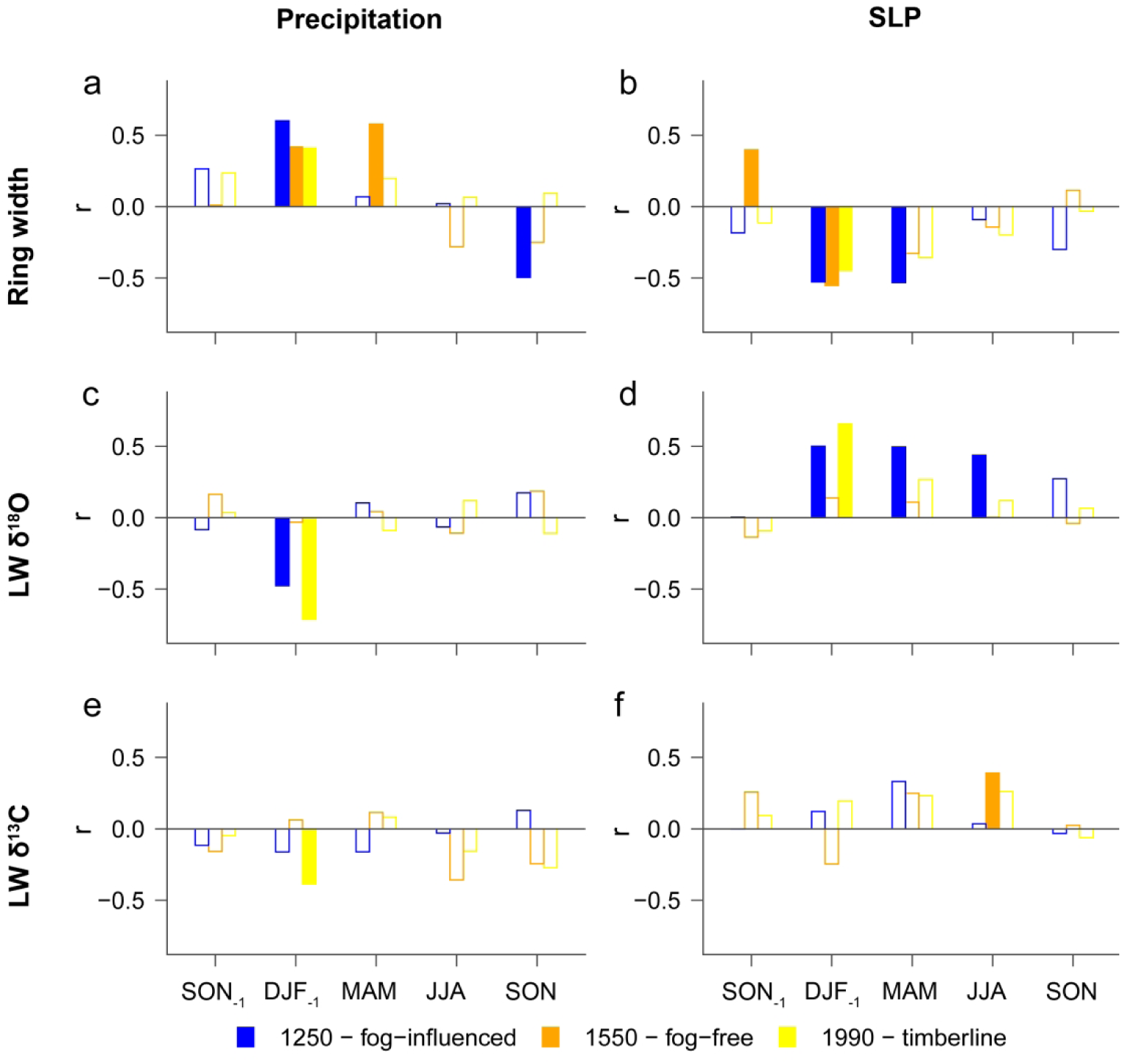
Statistical relationships between tree-ring variables and climate of the Canary Island pines studied on La Palma. Pearson’s correlation coefficients of ring widths and latewood oxygen and carbon isotope composition (LW δ^18^O, LW δ^13^C) of tree-ring cellulose of the studied period (2008-2017) from three sites (fog-influenced at 1250 m, fog-free at 1550 m and timberline at1990 m) with three-month averages of precipitation and sea-level pressure (SLP): previous growing season September, October, November (SON_-1_); December, January, February (DJF_-1_); current growing season March, April, May (MAM); June, July, August (JJA); September, October, November (SON). Filled bars represent significant correlations (P-value<0.05).

## Discussion

### δ^18^O variation in soil and plant tissues along the altitudinal gradient during the wet season

Soil water δ^18^O values decreased with increasing altitude (Fig. 2) as expected due to Rayleigh distillation known to occur for precipitation (Poage and Chamberlain 2001; Rozanski et al. 1993; Siegenthaler and Oeschger 1980). As no isotope fractionation is assumed to occur during root water uptake and transport in the xylem, δ^18^O values in stem and twig tissues followed this decrease. Stem and twig water in both xylem and phloem/bark tissues had similar isotope values, indicating water exchange between both tissues. The slight ^18^O enrichment in water from twig tissues at the highest site (1990 m) might be a consequence of mixing with ^18^O-enriched needle water and potentially enhanced twig evaporation due to lower relative humidity and/or higher VPD (Treydte et al. 2014). In contrast to soil and xylem water, δ^18^O values in needle water strongly increased along the altitudinal gradient. This is likely caused by the meteorological conditions at sampling time (a typical winter day with trade wind influence; Table 1,2), especially decreasing relative humidity and air temperature with increasing altitude. This was also predicted by the Craig-Gordon model for leaf isotope enrichment, where relative humidity and air temperature are important driving variables (Eq. 1). It further leads to the observed reduction of g*_s_* and transpiration rates, even if soil water availability was probably high due to recent precipitation events (Table 2). Canopy conductance in this species declines with increasing evaporative demand, especially in absence of soil drought conditions (Brito et al. 2014; Luis et al. 2005). Moreover, the increasing ^18^O enrichment in needle water along the altitudinal gradient was most probably imprinted on leaf assimilates (Lehmann et al. 2017). Our observation for the leaf level is supported by the altitudinal increase in δ^18^O values of tree-ring cellulose (Fig. 2 and Fig. 5). Finally, no differences were observed in δ^18^O values of water vapor along the altitudinal gradient during the fog-free sampling day. During foggy days, however, more variability may be found due to the temperature inversion towards higher altitudes preventing water vapor from rising up. This would in consequence also affect needle water δ^18^O values. In our case, we expect δ^18^O variations of tree-ring cellulose to be mainly modulated by the strong effect of evaporative enrichment, and further oxygen atom exchange with xylem water rather than by δ^18^O variation in water vapor. However, more detailed measurements of δ^18^O values of atmospheric water vapor during different meteorological conditions at different altitudes should be performed in future studies.

Besides the influence of environmental conditions, the observed variability of all studied tissues along the altitudinal gradient could also partly be caused by genetic variability (Prunier et al. 2016). Other studies on *P. canariensis*, however, emphasize that its genetic diversity and adaptations are mostly related to volcanism or extreme arid conditions at low south exposed slopes rather than on altitudinal gradients (López de Heredia et al. 2014; López et al. 2016; Navascués et al. 2006).

### Mechanistic modeling of needle water δ^18^O and the effect of needle anatomy

The general trend of increasing needle water ^18^O enrichment with decreasing humidity expected from the Craig-Gordon model was well reflected in our empiric data (despite the assumptions made for the 480 and 970 sites). However, modeled δ^18^O values of needle water at the 1800 m site were significantly lower than the observed δ^18^O values (P-value <0.05; Fig. 4). This result was consistent with the simulated leaf and air humidity simulations, and it could only be explained in the case of increased leaf temperatures in 2 standard deviations or with relative humidity values of −5 % or less below the measured conditions (Fig. S1). Even in those scenarios, values fell into the 1:1 line, that was also the case of the observed values elsewhere, meaning that the average oxygen composition of the whole needle water was similar to that at the evaporative site. This is untypical for conifer species as a thick-walled endodermis (Casparian strip) in needles is thought to isolate the source water of vascular bundle and transfusion tissue from evaporative enriched water (Roden et al. 2015; Timofeeva et al. 2020; Zwieniecki et al. 2007). Observed values might then be related to the progressive ^18^O enrichment (Fig. 3) that the exceptional long Canary Island pine needles (up to 30 cm length) showed, which is potentially driven by the diffusion of the heavy isotopologue from evaporative sites back to xylem water (through aquaporins), further enriching the longitudinal flow towards the tip (Cernusak et al. 2016; Helliker and Ehleringer 2002; Kannenberg et al. 2021). Differentiated transpiration rates along the needle surface could also trigger the longitudinal enrichment (Shu et al. 2008). Specific climatic or physiologic conditions (Table S1) might therefore play a role on discussed processes leading to a higher mixture of needle water and, thus, making isotope modeling particularly challenging in the extraordinary long Canary island pine needles.

The ^18^O enrichment along needle gradients was higher at 1250 m than at 1990 m (Fig. 3) and is likely explained by lower relative humidity at the mid-altitude site at the given sampling day (9^th^ of March), in contrast to inverse conditions of the previous day (7^th^ of March) due to changes in weather conditions (Table 2). The changes in relative humidity likely affected the needle water δ^18^O values more rapidly than the δ^18^O values of assimilates, so that the latter rather might reflect needle water δ^18^O values from previous days, which were clearly more ^18^O-enriched at the highest altitude (Fig. 2). This may also explain the unexpected opposite bottom-to-tip trends in the apparent biosynthetic oxygen isotope fractionation (εbio_app_) between needle water and Ne-WSC at both altitudes (Fig. 3). Nevertheless, the isotope trends in organic matter and tree-ring cellulose were similar (Fig. 2), which is important to know for the interpretation of the tree-ring isotope signals. Future studies on progressive ^18^O enrichment in needles in controlled conditions could decipher whether climate conditions (relative humidity), plant physiology (stomatal conductance), anatomy (Casparian strip and aquaporins) or a mix of these factors trigger the observed gradients.

### Foliar water uptake and impact of fog on tree-ring isotope patterns

On the Canaries, a distinct layer of fog typically develops during atmospheric stability and high atmospheric pressure in summer at mid altitude. We assessed for the first time the foliar water uptake capacity of the species (Table 4), implying an alternative way of water uptake that may relief water stress (e.g. summer drought) in its crown as already reported for other species (Burns Limm et al. 2009; Eller et al. 2013). However, further studies should be conducted to understand the mechanism that allows this process to occur.

Although fog-influenced and fog-free sites were not at the same altitude, we expected main differences to be caused by the contrast of fog presence/absence due to the slope orientation rather than other factors related to altitudinal difference (300 m). Higher tree-ring δ^18^O values of Canary Island pine at the fog-free (1550 m, WNW) than at the fog-influenced site (1250 m, NNE) are likely caused by lower stomatal conductance under lower relative humidity in fog-free conditions (Fig. 5, Table 1). Interestingly, fog water (with a closer geographic origin) was found to be ^18^O-enriched compared to advective precipitation during the wet season (with a further geographic origin) (Prada et al. 2015; Scholl et al. 2011). However, foggy environmental conditions (lower temperatures and irradiance, higher relative humidity and thus lower VPD) seemed to have a stronger effect on δ^18^O variations in needle water at mid-term than the ^18^O-enriched fog water input, leading to the impossibility to trace the δ^18^O of fog water in wood cellulose. The expected lower stomatal conductance at the fog-free site did not result in lower ^13^C-discrimination as tree-ring δ^13^C values were similar to the fog-influenced site. Likely, the fog-free site experienced also lower photosynthetic rates due to stomatal closure as previously observed in this species (Brito et al. 2014; Luis et al. 2005; Peters et al. 2003), causing similar intercellular CO_2_ concentrations and thus ^13^C discrimination at both sites. This would agree with Scheidegger’s et al (2000) model: Similar δ^13^C and higher δ^18^O values at the fog-free site would theoretically mean a reduction in both average maximum net photosynthesis and stomatal conductance (Table S2).

Tree rings at the timberline site (1990 m, NNE) were clearly enriched in both ^18^O and ^13^C compared to the fog-influenced and fog-free site (Fig. 5, Table 1). This site faces lower temperatures and drier air (i.e. lower relative humidity than the fog-free site, Table 1) and relatively low fog frequency and precipitation (similar to fog-free site). We therefore assume that low relative humidity caused low stomatal conductance (Table S2) and that this response was not compensated by the reduced photosynthetic rates, resulting in the highest δ^13^C and δ^18^O values of all sites in agreement with Scheidegger et al (2000).

### Seasonal isotopic variations and climate sensitivity of tree-ring parameters

The seasonal variation of δ^18^O at fog-free and timberline sites, with lower LW than EW δ^18^O values might be a result of changes in source water uptake from ^18^O-enriched (shallow) to ^18^O-depleted (deep) soil water, with the latter potentially originating from ^18^O-depleted winter precipitation (Allen et al. 2019; Allison et al. 1983; Goldsmith et al. 2012; Sarris et al. 2013; Treydte et al. 2014). This is also in agreement with the correlation of tree-ring width and LW δ^18^O with precipitation and SLP (Fig. 7). The role of winter precipitation on latewood formation is also supported by its negative correlation with LW δ^13^C, as higher soil water reserves might trigger a lower intrinsic water use efficiency in the pine needles and thus, a decrease in LW δ^13^C.

Seasonal δ^13^C differences were observed for tree rings at both the fog-influenced and fog-free sites (Fig. 5 b, d), with lower LW δ^13^C values compared to EW. At the fog-influenced site this could be caused by higher CO_2_ stomatal conductance and lower photosynthetic rates during the frequent foggy conditions in summer, thus more humid and with a lower irradiance (Marzol 2008; Ritter et al. 2019). Leaf gas exchange measurements during foggy conditions would need, however, to be performed to confirm this directly. At the fog-free site lower LW δ^13^C values compared to EW may be caused by the strong influence of stomatal closure on photosynthetic rates as discussed in the previous section and reported for this species (Brito et al. 2014; Luis et al. 2005; Peters et al. 2003; Scheidegger et al. 2000) (Table S2), with higher photosynthetic rates occurring during the winter days (Peters et al. 2008). Finally, lower temperatures, shorter growing seasons and generally lower photosynthetic activity at the timberline (Weigel et al. 2018) in addition to VPD conditions during latewood formation that reduced stomatal conductance and photosynthetic rate (Peters et al. 2003) (Table S2) may cause reduced intra-annual differences between mean EW and LW δ^13^C values. Thus, our results may primarily show that carbon and oxygen isotope patterns in tree rings of Canary Island pine reflect a strong interplay of plant physiology (stomatal regulation, water use efficiency) with environmental conditions (altitude-driven climate and seasonal variations).

EW and LW δ^18^O values converged at/in the more humid site/years whereas they diverged at/in drier sites/years. This is probably caused by lower humidity during LW formation. In correspondence to Kahmen et al. (2011) who explained 91 % of the variability of tree-ring cellulose δ^18^O with mean annual VPD, also a significant portion of the variability in EW and LW δ^18^O of our samples (68,5 % and 66.5 % respectively) was related to mean annual VPD. We, however, found mean annual relative humidity to have an even stronger effect on EW and LW δ^18^O (82.2 % and 78.5 % respectively) (Fig.6). This might be related to the different environmental conditions in both study regions with generally lower water availability and lower temperatures on the Canary Islands compared to Hawaii (Kahmen et al. 2011).

## Conclusions

Assessment of the carbon and oxygen isotope composition of various Canary Island pine tissues along an altitudinal gradient on La Palma showed the plasticity of physiological responses in this species to the highly variable environmental conditions along its broad climatic range of distribution. Our study demonstrates that changes in relative humidity along the altitudinal gradient exert the strongest influence on oxygen stable isotopes both at short (needle water) and midterm (tree-ring cellulose of 10 years). This, together with *in situ* measurements showed that the capacity of this species to adapt to the different studied environments is mainly triggered by the regulation of the water losses according to the local humidity conditions. The strong isotopic heterogeneity within the needle triggered by the relatively long needle morphology of the species, makes leaf water isotope modeling challenging. The similar δ^13^C values and thus water use efficiency during the wet and dry season at fog-free and fog-influenced sites show as well the adaptation of this pine to different environments. In addition, seasonal δ^18^O variations showed the ability of *P. canariensis* to uptake water from different soil water pools if needed. Moreover, we assessed for the first time the capability of this species to benefit from fog water by means of foliar uptake. However, fractionations derived from foggy environmental conditions left a greater imprint on cellulose δ^18^O than fog ^18^O-enriched water itself.

Plasticity of water loss regulation and water use efficiency of *P. canariensis* can partly explain its wide distribution across an archipelago with limited water input except at foggy sites. This broad distribution can also be a consequence of the little competition with other species on the Island and by the *P. canariensis* physiology, which is well-adapted to severe perturbances allowing it to prosper on volcanically active islands (López de Heredia et al. 2014; Miranda et al. 2020). The extreme physiological plasticity of *P. canariensis* could also allow the species to well-adapt to climatic changes in the near future.

## Acknowledgements

We thank Anne Verstege, Daniel Nievergelt, Frederick Reinig and Chiara Calderaro for great support in the field, and Manuela Oettli for the assistance in the isotope lab. We are grateful to the Cabildo and Forest Service of La Palma for logistic support. Climatic data was provided by the AEMET (Agencia Estatal de Meteorología).

## Conflict of interest

None declared.

## Funding

JCM was supported by the fellowship Emilio Gonzalez Esparcia and by the Ramon Areces Foundation. The study was funded by the Swiss National Science Foundation (SNF 200021_175888 ‘Troxy’ granted to KT, SNF Ambizione “TreeCarbo”, PZ00P2_179978, granted to MML). JA was supported by research grants 20-05840Y of the Czech Science Foundation, INTER-EXCELLENCE LTAUSA19137 provided by Czech Ministry of Education, Youth and Sports, and long-term research development project No. RVO 67985939 of the Czech Academy of Sciences.

## Supplementary data

**Figure S1:** Averaged observed vs. modeled δ^18^O values in needle water of Canary Island pine along an altitudinal gradient on La Palma when a) simulating a range of leaf temperatures above and below 2 standard deviations and b) simulating a range of relative humidity of −10 to +10 % changes. Black dotted line represents the 1:1 line. Empty circles represent significant differences (P-value <0.05) between simulated and observed values. Filled circles represent similar simulated and observed values.

**Figure S2:** Climate sensitivity of Canary Island pine tree rings on La Palma. Pearson’s correlation coefficients of ring widths, early and latewood oxygen and carbon isotope composition (EW δ^18^O, LW δ^18^O, EW δ^13^C, LW δ^13^C, respectively) of tree-ring cellulose of the studied period (2008-2017) from three sites (fog-influenced at 1250 m, fog-free at 1550 m and timberline at 1990 m) with three-month average of temperature, precipitation, sea-level pressure (SLP) and North Atlantic Oscillation (NAO): previous growing season September, October, November (SON_-1_); December, January, February (DJF_-1_); current growing season March, April, May (MAM); June, July, August (JJA); September, October, November (SON). Filled bars represent significant correlations (P<0.05).

**Table S1:** Mean input and output values of CG-model per site. T_air_: Air temperature, T_leaf_: leaf temperature, rH: relative humidity, g_s_: stomatal conductance, δ^18^O_WV_: water vapour δ^18^O, δ^18^O_TX_: twig xylem water δ^18^O, ε_eq_: equilibrium fractionation factor, ε_k_: kinetik fractionation factor, w_a_/w_i_: ratio of water vapor partial pressures outside and inside the leaf, δ^18^O_Ne-Wat_: needle water δ^18^O.

**Table S2:** Average number of days per month in which a vapor pressure deficit > 2.5 kPa is reached. According to Peters Morales and Jiménez (2003), stomatal conductance decreases dramatically above the 2.5 kPa threshold, reaching zero conductance at a VPD of 4.5 kPa.

## Notes

### Competing Interest Statement

The authors have declared no competing interest.

## References

Aboal JR, Jimenez MS, Morales D, Gil P (2000) Effects of thinning on throughfall in Canary Islands pine forest - the role of fog. Journal of Hydrology. 238:218–230.

Aboal JR, Regalado C, Ritter A, Gómez LA, Fernández AB (2013) Interceptación de lluvia y niebla en bosques de laurisilva y pinar de las Islas Canarias. In: Cabezas F (ed) Interceptación de la lluvia por la vegetación en España. Instituto Euromediterráneo del Agua, Murcia, Spain, pp 47–78.

AEMET-IM (2012) Atlas climático de los archipiélagos de Canarias, Madeira y Azores. Agencia estatal de meteorología (MAGAMA) e Instituto de meteorologia de Portugal.

Aguilar MJA, Gonzalez-Gonzalez R, Garzon-Machado V, Pizarro-Hernandez B (2010) Actual and potential natural vegetation on the Canary Islands and its conservation status. Biodiversity and Conservation. 19:3089–3140.

Allen ST, Kirchner JW, Braun S, Siegwolf RTW, Goldsmith GR (2019) Seasonal origins of soil water used by trees. Hydrology and Earth System Sciences. 23:1199–1210.

Allison GB, Barnes CJ, Hughes MW (1983) THE DISTRIBUTION OF DEUTERIUM AND O-18 IN DRY SOILS .2. EXPERIMENTAL. Journal of Hydrology. 64:377–397.

Barbour MM (2007) Stable oxygen isotope composition of plant tissue: a review. Functional Plant Biology. 34:83–94.

Bechtel B (2016) The climate of the Canary Islands by annual cycle parameters. International Archives of the Photogrammetry Remote Sensing and Spatial Information Sciences. 41:243–250.

Boegelein R, Lehmann MM, Thomas FM (2019) Differences in carbon isotope leaf-to-phloem fractionation and mixing patterns along a vertical gradient in mature European beech and Douglas fir. New Phytologist. 222:1803–1815.

Boettger T, Haupt M, Knoeller K, Weise SM, Waterhouse JS, Rinne KT, Loader NJ, Sonninen E, Jungner H, Masson-Delmotte V, Stievenard M, Guillemin M-T, Pierre M, Pazdur A, Leuenberger M, Filot M, Saurer M, Reynolds CE, Helle G, Schleser GH (2007) Wood cellulose preparation methods and mass spectrometric analyses of delta C-13, delta O-18, and nonexchangeable delta H-2 values in cellulose, sugar, and starch: An interlaboratory comparison. Analytical Chemistry. 79:4603–4612.

Brinkmann N, Seeger S, Weiler M, Buchmann N, Eugster W, Kahmen A (2018) Employing stable isotopes to determine the residence times of soil water and the temporal origin of water taken up by Fagus sylvatica and Picea abies in a temperate forest. New Phytologist. 219:1300–1313.

Brito P, Lorenzo JR, Gonzalez-Rodriguez AM, Morales D, Wieser G, Jimenez MS (2014) Canopy transpiration of a Pinus canariensis forest at the tree line: implications for its distribution under predicted climate warming. European Journal of Forest Research. 133:491–500.

Brueggemann N, Gessler A, Kayler Z, Keel SG, Badeck F, Barthel M, Boeckx P, Buchmann N, Brugnoli E, Esperschuetz J, Gavrichkova O, Ghashghaie J, Gomez-Casanovas N, Keitel C, Knohl A, Kuptz D, Palacio S, Salmon Y, Uchida Y, Bahn M (2011) Carbon allocation and carbon isotope fluxes in the plant-soil-atmosphere continuum: a review. Biogeosciences. 8:3457–3489.

Burgess SSO, Dawson TE (2004) The contribution of fog to the water relations of Sequoia sempervirens (D. Don): foliar uptake and prevention of dehydration. Plant Cell and Environment. 27:1023–1034.

Burk RL, Stuiver M (1981) OXYGEN ISOTOPE RATIOS IN TREES REFLECT MEAN ANNUAL TEMPERATURE AND HUMIDITY. Science. 211:1417–1419.

Burns Limm E, Simonin KA, Bothman AG, Dawson TE (2009) Foliar water uptake: a common water acquisition strategy for plants of the redwood forest. Oecologia. 161:449–459.

Cernusak LA, Barbour MM, Arndt SK, Cheesman AW, English NB, Feild TS, Helliker BR, Holloway-Phillips MM, Holtum JAM, Kahmen A, McInerney FA, Munksgaard NC, Simonin KA, Song X, Stuart-Williams H, West JB, Farquhar GD (2016) Stable isotopes in leaf water of terrestrial plants. Plant Cell and Environment. 39:1087–1102.

Climent J, Gil L, Tuero M (1996) Las regiones de procedencia de Pinus canariensis Chr Sm. ex DC. ICONA, Madrid.

Craig H, Gordon LI. 1965. Deuterium and oxygen-18 variations in the ocean and the marine atmosphere. In Stable Isotopes in Oceanographic Studies and Paleotemperatures Ed. Tongiorgi E, Spoleto, Italy: Lischi and Figli, pp 9–130.

Dorta P (1985) La inversión térmica en Canarias. Investigaciones Geográficas. 15:109–126.

Eidg. Forschungsanstalt WSL. Pines in the Mist - Fast Forward Science 2018. https://www.youtube.com/watch?v=wBmOWmTrglw (20 November 2020 date last accessed|)|.

Eller CB, Lima AL, Oliveira RS (2013) Foliar uptake of fog water and transport belowground alleviates drought effects in the cloud forest tree species, Drimys brasiliensis (Winteraceae). New Phytologist. 199:151–162.

Farquhar GD, Cernusak LA, Barnes B (2007) Heavy water fractionation during transpiration. Plant Physiology. 143:11–18.

Gartner H, Nievergelt D (2010) The core-microtome: A new tool for surface preparation on cores and time series analysis of varying cell parameters. Dendrochronologia. 28:85–92.

Gehre M, Geilmann H, Richter J, Werner RA, Brand WA (2004) Continuous flow H-2/H-1 and and(18)O/O-16 analysis of water samples with dual inlet precision. Rapid Communications in Mass Spectrometry. 18:2650–2660.

Gessler A, Pedro Ferrio J, Hommel R, Treydte K, Werner RA, Monson RK (2014) Stable isotopes in tree rings: towards a mechanistic understanding of isotope fractionation and mixing processes from the leaves to the wood. Tree Physiology. 34:796–818.

Gessler A, Treydte K (2016) The fate and age of carbon - insights into the storage and remobilization dynamics in trees. New Phytologist. 209:1338–1340.

Goldsmith GR, Matzke NJ, Dawson TE (2013) The incidence and implications of clouds for cloud forest plant water relations. Ecology Letters. 16:307–314.

Goldsmith GR, Munoz-Villers LE, Holwerda F, McDonnell JJ, Asbjornsen H, Dawson TE (2012) Stable isotopes reveal linkages among ecohydrological processes in a seasonally dry tropical montane cloud forest. Ecohydrology. 5:779–790.

Harris I, Jones PD, Osborn TJ, Lister DH (2014) Updated high-resolution grids of monthly climatic observations - the CRU TS3.10 Dataset. International Journal of Climatology. 34:623–642.

Helle G, Schleser GH (2004) Beyond CO2-fixation by Rubisco - an interpretation of C-13/C-12 variations in tree rings from novel intra-seasonal studies on broad-leaf trees. Plant Cell and Environment. 27:367–380.

Helliker BR, Ehleringer JR (2002) Differential O-18 enrichment of leaf cellulose in C-3 versus C-4 grasses. Functional Plant Biology. 29:435–442.

Herrera RG, Puyol DG, Martin EH, Gimeno L, Rodriguez PR (2001) Influence of the North Atlantic Oscillation on the Canary Islands precipitation. Journal of Climate. 14:3889–3903.

Holmes R (1983) Computer Assisted Quality Control in Tree-Ring Dating and Measurement. Tree-Ring Bulletin. 44:69–75.

Hu YS, Yao BJ (1981) TRANSFUSION TISSUE IN GYMNOSPERM LEAVES. Botanical Journal of the Linnean Society. 83:263–272.

Janecka K, Kaczka RJ, Gartner H, Harvey JE, Treydte K (2020) Compression wood has a minor effect on the climate signal in tree-ring stable isotope records of montane Norway spruce. Tree Physiology. 40:1014–1028.

Johnstone JA, Roden JS, Dawson TE (2013) Oxygen and carbon stable isotopes in coast redwood tree rings respond to spring and summer climate signals. Journal of Geophysical Research-Biogeosciences. 118:1438–1450.

Kahmen A, Sachse D, Arndt SK, Tu KP, Farrington H, Vitousek PM, Dawson TE (2011) Cellulose delta O-18 is an index of leaf-to-air vapor pressure difference (VPD) in tropical plants. Proceedings of the National Academy of Sciences of the United States of America. 108:1981–1986.

Kannenberg SA, Fiorella RP, Anderegg WRL, Monson RK, Ehleringer JR (2021) Seasonal and diurnal trends in progressive isotope enrichment along needles in two pine species. Plant Cell and Environment. 44:143–155.

Kim K, Lee X (2011) Transition of stable isotope ratios of leaf water under simulated dew formation. Plant Cell and Environment. 34:1790–1801.

Körner C, Farquhar GD, Wong SC (1991) Carbon isotope discrimination by plants follows latitudinal and altitudinal trends. Oecologia. 88:30–40.

Laumer W, Andreu L, Helle G, Schleser GH, Wieloch T, Wissel H (2009) A novel approach for the homogenization of cellulose to use micro-amounts for stable isotope analyses. Rapid Communications in Mass Spectrometry. 23:1934–1940.

Lehmann MM, Gamarra B, Kahmen A, Siegwolf RTW, Saurer M (2017) Oxygen isotope fractionations across individual leaf carbohydrates in grass and tree species. Plant Cell and Environment. 40:1658–1670.

Lehmann MM, Rinne KT, Blessing C, Siegwolf RTW, Buchmann N, Werner RA (2015) Malate as a key carbon source of leaf dark-respired CO2 across different environmental conditions in potato plants. Journal of Experimental Botany. 66:5769–5781.

Limm EB, Simonin KA, Bothman AG, Dawson TE (2009) Foliar water uptake: a common water acquisition strategy for plants of the redwood forest. Oecologia. 161:449–459.

López de Heredia U, López R, Collada C, Emerson BC, Gil L (2014) Signatures of volcanism and aridity in the evolution of an insular pine (Pinus canariensis Chr. Sm. Ex DC in Buch). Heredity. 113:240–249.

López R, Cano FJ, Choat B, Cochard H, Gil L (2016) Plasticity in vulnerability to cavitation of Pinus canariensis occurs only at the driest end of an aridity gradient. Frontiers in Plant Science. 7

López R, Climent J, Gil L (2008) From desert to cloud forest: the non-trivial phenotypic variation of Canary Island pine needles. Trees-Structure and Function. 22:843–849.

López R, Climent J, Gil L (2010) Intraspecific variation and plasticity in growth and foliar morphology along a climate gradient in the Canary Island pine. Trees-Structure and Function. 24:343–350.

López R, López de Heredia U, Collada C, Javier Cano F, Emerson BC, Cochard H, Gil L (2013) Vulnerability to cavitation, hydraulic efficiency, growth and survival in an insular pine (Pinus canariensis). Annals of Botany. 111:1167–1179.

Luis VC, Jimenez MS, Morales D, Kucera J, Wieser G (2005) Canopy transpiration of a Canary islands pine forest. Agricultural and Forest Meteorology. 135:117–123.

Luis VC, Taschler D, Hacker J, Jimenez MS, Wieser G, Neuner G (2007) Ice nucleation and frost resistance of Pinus canariensis seedlings bearing needles in three different developmental states. Annals of Forest Science. 64:177–182.

Marzol MV (2008) Temporal characteristics and fog water collection during summer in Tenerife (Canary Islands, Spain). Atmospheric Research. 87:352–361.

Marzol MV, Máyer P (2012) Algunas reflexiones acerca del clima de las Islas Canarias. Nimbus. 29–30:399-416.

Máyer P, Marzol MV. 2013. Análisis de la pluviosidad en las islas canarias mediante la elaboración de gradientes. *In* XIII Congreso de Geógrafos Españoles. Universitat de les Illes Balears, Spain, pp 145–154.

McCarroll D, Loader NJ (2004) Stable isotopes in tree rings. Quaternary Science Reviews. 23:771–801.

Miranda JC. 2017. El pino canario y las erupciones de Cumbre Vieja (1949, La Palma).Adaptación al volcanismo. Universidad Politécnica de Madrid, Madrid, p 127.

Miranda JC, Rodríguez-Calcerrada J, Pita P, Saurer M, Oleksyn J, Gil L (2020) Carbohydrate dynamics in a resprouting species after severe aboveground perturbations. European Journal of Forest Research

Naoe S, Tayasu I, Masaki T, Koike S (2016) Negative correlation between altitudes and oxygen isotope ratios of seeds: exploring its applicability to assess vertical seed dispersal. Ecology and Evolution. 6:6817–6823.

Navascués M, Vaxevanidou Z, Gonzalez-Martinez SC, Climent J, Gil L, Emerson BC (2006) Chloroplast microsatellites reveal colonization and metapopulation dynamics in the Canary Island pine. Molecular Ecology. 15:2691–2698.

Offermann C, Pedro Ferrio J, Holst J, Grote R, Siegwolf R, Kayler Z, Gessler A (2011) The long way down-are carbon and oxygen isotope signals in the tree ring uncoupled from canopy physiological processes? Tree Physiology. 31:1088–1102.

Pérez-de-Paz P, Salas M, Rodríguez O, Acebes J, Del-Arco M, Wildpret W (1994) Atlas Cartográfico de los Pinares Canarios IV: Gran Canaria y plantaciones de Fuerteventura y Lanzarote. Viceconsejería de Medio Ambiente, Consejería de Política Territorial, Gobierno de Canarias, Santa Cruz de Tenerife, Spain.

Peters J, Gonzalez-Rodriguez AM, Jimenez MS, Morales D, Wieser G (2008) Influence of canopy position, needle age and season on the foliar gas exchange of Pinus canariensis. European Journal of Forest Research. 127:293–299.

Peters J, Morales D, Jimenez MS (2003) Gas exchange characteristics of Pinus canariensis needles in a forest stand on Tenerife, Canary Islands. Trees-Structure and Function. 17:492–500.

Pinheiro J, Bates D, DebRoy D, Sarkar D, Team RC. {nlme}: Linear and Nonlinear Mixed Effects Models. https://CRAN.R-project.org/package=nlme (20 November 2020 date last accessed|)|.

Poage MA, Chamberlain CP (2001) Empirical relationships between elevation and the stable isotope composition of precipitation and surface waters: Considerations for studies of paleoelevation change. American Journal of Science. 301:1–15.

Prada S, Figueira C, Aguiar N, Cruz JV (2015) Stable isotopes in rain and cloud water in Madeira: contribution for the hydrogeologic framework of a volcanic island. Environmental Earth Sciences. 73:2733–2747.

Prunier J, Verta JP, MacKay JJ (2016) Conifer genomics and adaptation: at the crossroads of genetic diversity and genome function. New Phytologist. 209:44–62.

R Core Team. 2019. R: A language and environment for statistical computing. R Foundation for Statistical Computing, Viena.

Ritter A, Regalado CM, Aschan G (2008) Fog Water Collection in a Subtropical Elfin Laurel Forest of the Garajonay National Park (Canary Islands): A Combined Approach Using Artificial Fog Catchers and a Physically Based Impaction Model. Journal of Hydrometeorology. 9:920–935.

Ritter A, Regalado CM, Guerra JC (2019) The impact of climate change on water fluxes in a Macaronesian cloud forest. Hydrological Processes. 33:2828–2846.

Roden J, Kahmen A, Buchmann N, Siegwolf R (2015) The enigma of effective path length for O-18 enrichment in leaf water of conifers. Plant Cell and Environment. 38:2551–2565.

Rodríguez Martín JA, Nanos N, Miranda JC, Carbonell G, Gil L (2013) Volcanic mercury in Pinus canariensis. Naturwissenschaften. 100:739–747.

Rozanski K, Araguás-Araguás L, Gonfiantini R (1993) Isotopic Patterns in Modern Global Precipitation. In: Swart PK, Lohmann KC, McKenzie J, Savin S (eds) Climate Change in Continental Isotopic Records. American Geophysical Union, Washington DC, pp 1–36.

Rozas V, Garcia-Gonzalez I, Perez-de-Lis G, Arevalo JR (2013) Local and large-scale climatic factors controlling tree-ring growth of Pinus canariensis on an oceanic island. Climate Research. 56:197–207.

Sarris D, Siegwolf R, Koerner C (2013) Inter- and intra-annual stable carbon and oxygen isotope signals in response to drought in Mediterranean pines. Agricultural and Forest Meteorology. 168:59–68.

Scheidegger Y, Saurer M, Bahn M, Siegwolf R (2000) Linking stable oxygen and carbon isotopes with stomatal conductance and photosynthetic capacity: a conceptual model. Oecologia. 125:350–357.

Scholl M, Eugster W, Burkard R (2011) Understanding the role of fog in forest hydrology: stable isotopes as tools for determining input and partitioning of cloud water in montane forests. Hydrological Processes. 25:353–366.

Shu Y, Feng XH, Posmentier ES, Sonder LJ, Faiia AM, Yakir D (2008) Isotopic studies of leaf water. Part 1: A physically based two-dimensional model for pine needles. Geochimica Et Cosmochimica Acta. 72:5175–5188.

Shuttleworth WJ (1977) The exchange of wind-driven fog and mist between vegetation and the atmosphere. Boundary-Layer Meteorology. 12:463–489.

Siegenthaler U, Oeschger H (1980) Correlation of O-18 in precipitation with temperature and altitude. Nature. 285:314–317.

Sternberg L, Deniro MJD (1983) Biogeochemical implications of the isotopic equilibrium fractionation factor between the oxygen-atoms of acetone and water. Geochimica Et Cosmochimica Acta. 47:2271–2274.

Sternberg L, Pinzon MC, Anderson WT, Jahren AH (2006) Variation in oxygen isotope fractionation during cellulose synthesis: intramolecular and biosynthetic effects. Plant Cell and Environment. 29:1881–1889.

Stokes MA, Smiley TL (1996) An Introduction to Tree-ring Dating. University of Arizona Press, Chicago, IL, USA.

Tang KL, Feng XH (2001) The effect of soil hydrology on the oxygen and hydrogen isotopic compositions of plants’ source water. Earth and Planetary Science Letters. 185:355–367.

Terwilliger VJ, Betancourt JL, Leavitt SW, Van de Water PK (2002) Leaf cellulose delta D and delta O-18 trends with elevation differ in direction among co-occurring, semiarid plant species. Geochimica Et Cosmochimica Acta. 66:3887–3900.

Timofeeva G, Treydte K, Bugmann H, Salmon Y, Rigling A, Schaub M, Vollenweider P, Siegwolf R, Saurer M (2020) How does varying water supply affect oxygen isotope variations in needles and tree rings of Scots pine? Tree Physiology. 40:1366–1380.

Torres C, Cuevas E, Guerra J, Carreno V (2001) Caracterización de las masas de aire en la región subtropical Libro de comunicaciones del V Simposio Nacional de Predicción. Instituto Nacional de Meteorología, Madrid.

Treydte K, Boda S, Pannatier EG, Fonti P, Frank D, Ullrich B, Saurer M, Siegwolf R, Battipaglia G, Werner W, Gessler A (2014) Seasonal transfer of oxygen isotopes from precipitation and soil to the tree ring: source water versus needle water enrichment. New Phytologist. 202:772–783.

Valladares P. 1995. Estudio geográfico del mar de nubes en la vertiente norte de Tenerife. *In* Departamento de Geografía. Universidad de la Laguna, Tenerife.

Weigel R, Irl SDH, Treydte K, Beierkuhnlein C, Berels J, Field R, Miranda JC, Steinbauer A, Steinbauer MJ, Jentsch A (2018) A novel dendroecological method finds a non-linear relationship between elevation and seasonal growth continuity on an island with trade wind-influenced water availability. Aob Plants. 10

West AG, Patrickson SJ, Ehleringer JR (2006) Water extraction times for plant and soil materials used in stable isotope analysis. Rapid Communications in Mass Spectrometry. 20:1317–1321.

Wickham H (2009) ggplot2: Elegant Graphics for Data Analysis. Ggplot2: Elegant Graphics for Data Analysis:1–212.

Zwieniecki MA, Brodribb TJ, Holbrook NM (2007) Hydraulic design of leaves: insights from rehydration kinetics. Plant Cell and Environment. 30:910–921.

